# Methylation of histone H3 lysine 4 is required for maintenance of beta cell function in adult mice

**DOI:** 10.1101/2021.01.28.428651

**Authors:** Ben Vanderkruk, Nina Maeshima, Daniel J. Pasula, Meilin An, Cassandra L. McDonald, Francis C. Lynn, Dan S. Luciani, Brad G. Hoffman

**Affiliations:** Diabetes Research Group, BC Children’s Hospital Research Institute, Vancouver, BC, Canada; Department of Surgery, University of British Columbia, Vancouver, BC, Canada

**Keywords:** H3K4me3, DPY30, COMPASS, Transcription, Beta cell, Type 2 diabetes

## Abstract

Pancreatic β-cells control glucose homeostasis via regulated production and secretion of insulin. This function arises from a highly specialized gene expression program which is established during development and then sustained, with limited flexibility, in terminally differentiated β-cells. Dysregulation of this program is seen in type 2 diabetes (T2D) but mechanisms that preserve gene expression or underlie its dysregulation in mature β-cells are not well resolved. Here we show that trithorax group-dependent histone H3 lysine 4 trimethylation (H3K4me3) maintains expression of genes important for insulin biosynthesis and glucose-responsiveness in β-cells. Transcriptional changes in H3K4me3-deficient β-cells lead to severe hyperglycemia in adult mice. We show that H3K4me3 deficiency leads to a less active and more repressed epigenome profile, which locally correlates with gene expression deficits but does not globally reduce gene expression. Instead, developmentally regulated genes and genes in weakly active or suppressed states particularly rely on H3K4 methylation. We then show that H3K4me3 is re-organized in diabetic *Lepr^db/db^* mouse islets in favour of weakly active and disallowed genes at the expense of terminal β-cell markers with broad H3K4me3 peaks. Our results point to key roles of H3K4me3 in maintaining mature β-cell function and establishing a dysfunctional transcriptome in diabetic islets.

## Introduction

Pancreatic islets are endocrine micro-organs that regulate glucose homeostasis. In adults, islet β-cells are the exclusive source of circulating insulin, the major blood glucose-lowering hormone. Insufficient insulin release by β-cells is the primary etiology of type 2 diabetes (T2D). The rising global incidence of T2D ^1^ has prompted intensive research into mechanisms underlying the progressive dysfunction of β-cells that leads to T2D. Transcriptome remodeling, involving activation of developmentally silenced genes and decreased expression of genes important for insulin biosynthesis, glucose-responsiveness, or those that enforce the β-cell identity, emerges in β-cells exposed to chronic metabolic stress ^2^. While β-cell-enriched transcription factors have been the primary focus of studies defining the transcriptional program in mature β-cells, there is mounting evidence that chromatin modifications play a central role in the maintenance of mature β-cell identity and function, and that the chromatin landscape is reshaped in β- cells exposed to chronic metabolic stress and T2D ^3–10^.

Tissue-specific transcription programs rely on precise activation and maintenance of specific genes and stable repression of other genes. Expression patterns are established during cell differentiation and maturation, and then maintained during the balance of that cell’s life, which may span decades, cell divisions, and environmental perturbations. Active and repressed genes have stereotypical patterns of chromatin modification which may stabilize their transcription state and are useful in their systematic classification ^11^. For example, it has been shown that H3K4me3 is reliably enriched on nucleosomes at the 5’ end of actively transcribed genes and is therefore the canonical marker of active promoters ^12^. Despite its utility as a marker of promoter activity, global (i.e., genome-wide) decreases in H3K4 methylation are remarkably well tolerated in embryonic stem cells and during development in genetic knockout models ^13–16^. Evidence in multicellular animal models points to a requirement for H3K4 methyltransferase activities during certain specialized developmental events, whereas evidence for continued requirement in mature tissues is rare ^16–21^.

Mono-, di-, and trimethylation of H3K4 in mammalian cells is predominantly catalyzed by COMPASS and COMPASS-related complexes. They are composed of the core subunits WDR5, RBBP5, ASH2L, and DPY30, plus one of six catalytic subunits MLL1-4 or SET1A-B, and a variable retinue of accessory factors ^22^ . We previously defined roles for DPY30 during embryonic pancreas development. DPY30 fine-tunes cell fate decisions to decrease endocrine acinar cell specification in favour of exocrine cell specification ^21^ and helps to activate terminal marker genes during endocrine cell maturation ^23^. By contrast, it is not known whether the continuous activity of H3K4 methyltransferases is necessary in terminally differentiated islet cells, where lineage-specific transcription programs have already been established. In islets, recruitment of H3K4 methyltransferase proteins by transcription factors—for example, recruitment of SETD7 by PDX1 ^7^, and MLL3/4 by MAFA ^24^—support expression of insulin and other genes that are important for mature β-cell functions. Whether these functions are performed by H3K4 methylation or by independent coactivator activities of these complexes is not clear ^14, 21, 25^. We therefore sought to define the role for H3K4 methylation in the regulation of gene expression in mature β-cells.

## Results

### Reduction to H3K4 methylation in beta cells of adult mice leads to glucose intolerance and hyperglycemia

With the aim of reducing H3K4 methylation we targeted DPY30, a noncatalytic core subunit of all COMPASS complexes which is not required for complex assembly or coactivator activity but boosts methyltransferase activities ^26, 27^. To induce synchronized deletion of *Dpy30* in mature β-cells, mice harbouring a Pdx1:*CreER^TM^* transgene ^28^ and *LoxP* sites flanking the fourth exon of *Dpy30* ^21^ were administered tamoxifen at eight weeks of age (herein called *Dpy30*-KO) (Fig 1A). As expected, knockout of *Dpy30* did not impact assembly of the other core subunits of the COMPASS complex WDR5, RBBP5, and ASH2L, or their association with chromatin (Fig 1B) but did lead to reductions of H3K4me3 and H3K4me1 in islets (Fig 1C).

**Figure 1.**
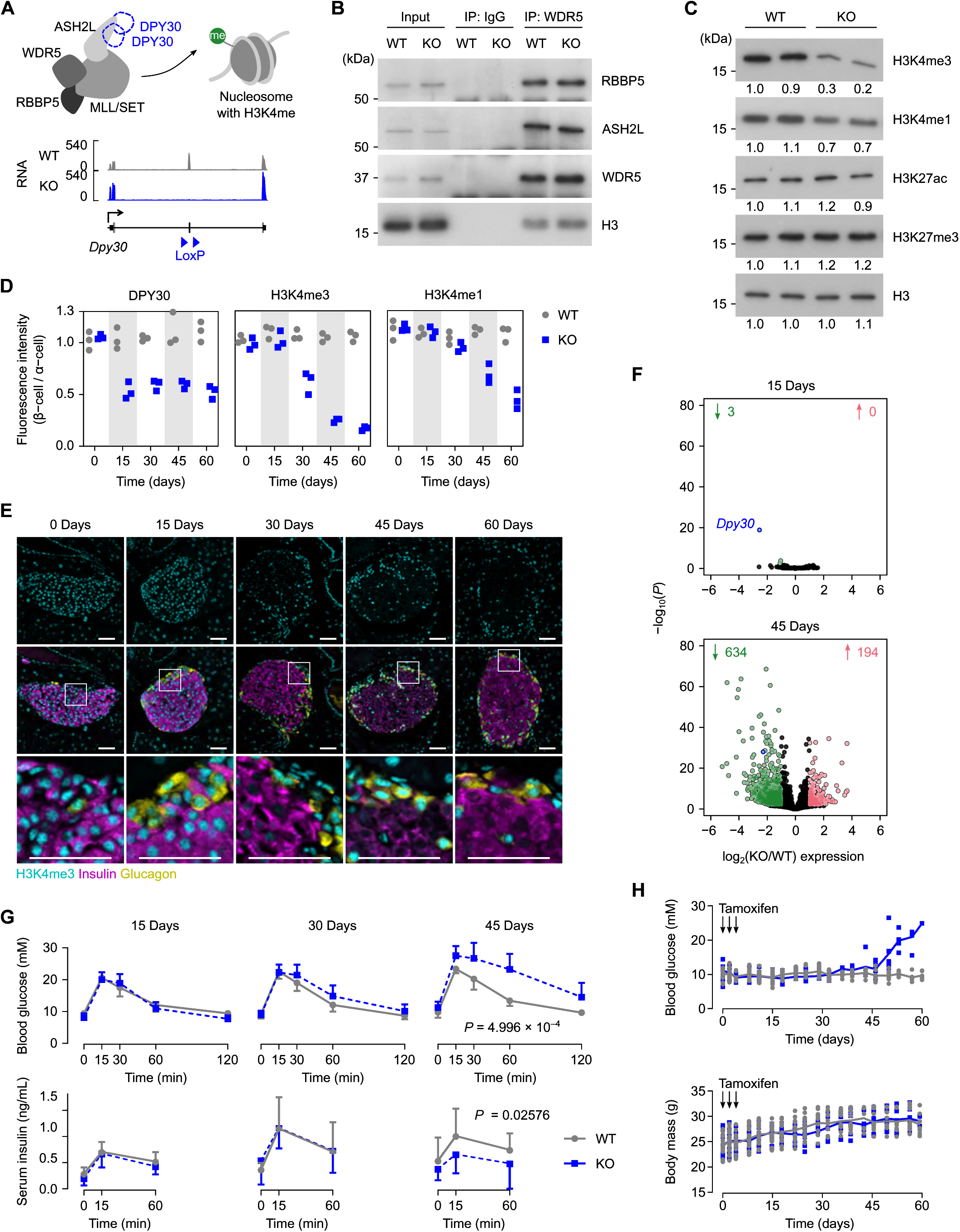
Reduction to H3K4 methylation in beta cells leads to glucose intolerance and hyperglycemia in adult mice. **A,** Top: schematic showing the core subunits of COMPASS complexes and a nucleosome methylated on H3K4. Bottom: genome-aligned RNA-seq reads at the *Dpy30* gene locus in *Dpy30*-WT and -KO cells. **B,** Immunoblots showing COMPASS core subunits RBBP5, ASH2L, and WDR5, and nucleosome protein histone H3, coimmunoprecipitated with WDR5 or an IgG control from *Dpy30*-WT or -KO islet cell nuclei 45 days after tamoxifen administration. Representative immunoblots of three independent coIPs are shown. IP: immunoprecipitation. **C,** Immunoblots showing H3K4me3, H3K4me1, H3K27ac, H3K27me3, and total histone H3 in islets from *Dpy30*-WT and -KO mice 45 days after tamoxifen administration. Numbers beneath each band indicate the band intensity normalized to the left-most sample. **D,** Mean immunofluorescent intensity of DPY30, H3K4me3, and H3K4me1 in *Dpy30*-WT and -KO β-cell nuclei at the indicated days after tamoxifen administration. Data are normalized to fluorescence intensity of α-cell nuclei; n = 3 mice of each genotype and time-point. **E,** Example IHC images of *Dpy30*-KO islets used for measurements in panel D showing H3K4me3 (cyan), insulin (magenta), and glucagon (yellow). Scale bars: 100 μm. **F,** Scatterplots showing log2(fold-change) and -log10(*P*-value) of gene expression in *Dpy30*-KO versus -WT cells 15- and 45-days after tamoxifen administration. Genes showing a ≥2-fold increase or decrease in expression with *P* ≤ 0.01 (Wald test with Benjamini-Hochberg correction) are coloured red or green, respectively, and enumerated above. The full-length transcript of *Dpy30* is outlined and labeled in blue. **G,** Blood glucose (top) and serum insulin (bottom) concentration during IPGTT in *Dpy30*-WT and -KO mice 15-, 30-, and 45-days after tamoxifen administration. Data presented as mean ± SD (n = 8 mice for glucose measurements, 8-15 mice for insulin measurements). *P*-values were calculated from area under the curve (AUC) using multiple two-tailed t-tests with Welch’s and Benjamini-Hochberg corrections; only values ≤ 0.05 are shown. **H,** Unfasted blood glucose (top) and body mass (bottom) of *Dpy30*-WT and -KO mice during the 60-days after tamoxifen administration. Data are presented as individual measurements with mean (n = 8, however tracking was stopped after a blood glucose reading ≥ 20 mM).

We tracked the loss of H3K4 methylation in *Dpy30*-KO β-cells using immunofluorescence at 15- day intervals after tamoxifen administration. As seen in Fig 1D-E, H3K4me3 was stable for at least 15 days and then gradually lost during the subsequent 45 days. H3K4me1 showed a similar pattern, but its loss was even more gradual, whereas DPY30 was undetectable by 15 days after tamoxifen administration (Fig 1D). We took advantage of the slow turnover of H3K4me3 and H3K4me1 to parse the relative contribution of H3K4 methylation, versus DPY30 itself, in controlling gene expression in mature β-cells. We reasoned that defects arising in *Dpy30*-KO β-cells by 15-days post-tamoxifen (when DPY30 is lost but H3K4 methylation is unchanged) highlight functions of DPY30 itself. Defects arising later, by 45-days post-tamoxifen (when H3K4 methylation is reduced) highlight functional contributions of H3K4 methylation. To measure the β-cell transcriptome, GFP-positive islet cells labeled with the mTmG reporter ^29^ were enriched using fluorescence-activated cell sorting (FACS) from *Dpy30*-KO and -WT mice 15- and 45-days post-tamoxifen and processed for calibrated bulk RNA-seq analysis ^30^. Remarkably, loss of DPY30 had little effect on gene expression in β-cells, with only three genes (*Dpy30*, *Edn3*, *C3*) identified as differentially expressed 15-days post-tamoxifen (Fig 1F). In contrast, 828 genes were dysregulated 45-days post-tamoxifen, the majority of which (634) showed lower expression in *Dpy30*-KO cells (Fig 1F). We conclude that transcriptome remodeling in *Dpy30*-KO β-cells results from loss of H3K4 methylation.

To determine whether loss of DPY30 and/or H3K4 methylation in β-cells impairs glucose homeostasis in adult mice, we measured glucose clearance and insulin levels after intraperitoneal injection of glucose (2 g/kg body weight) in mice that had been fasted for 6 hrs. While glucose tolerance was no different between *Dpy30*-WT and -KO mice 15- and 30-days after tamoxifen administration, *Dpy30*-KO mice developed impaired blood glucose clearance and reduced serum insulin levels 45-days post-tamoxifen (Fig 1G). Glucose intolerance quickly progressed to hyperglycemia (Fig 1H) prompting euthanasia by 60-days.

Since the Pdx1:*CreER^TM^* transgene is known to also drive recombination in pancreatic islet δ-cells ^31^ and hypothalamic neurons ^32^ we additionally used the *Ins1^Cre^* mouse model ^33^ to confirm that *Dpy30* deletion specifically in β-cells is sufficient to drive the *in vivo* phenotype. Deletion of *Dpy30* from maturing β-cells using *Ins1^Cre^* caused reduction of H3K4me3 and H3K4me1 in islets by 5-weeks of age (Fig S1A-B). Unfasted blood glucose was dramatically elevated from 4- to 5-weeks of age, whereas body mass was similar among *Dpy30*-KO, -WT, and -heterozygous *Ins1^Cre^* mice (Fig S1C-D). *Ins1^Cre^ Dpy30*-KO mice showed impaired glucose clearance and low serum insulin following intraperitoneal glucose injection (Fig S1E-F), confirming that β-cell-intrinsic dysfunction is sufficient to drive metabolic defects in *Dpy30*-KO mice. In summary, reduction of H3K4 methylation in β-cells leads to impaired glucose tolerance, reduced serum insulin, and hyperglycemia.

### H3K4me3 maintains expression of genes involved in insulin production and glucose-stimulated activity

The reduction of serum insulin in *Dpy30*-KO mice prompted us to examine insulin production and secretion. We first confirmed that H3K4me3 was lost from the *Ins1* and *Ins2* promoters in Pdx1:*CreER^TM^ Dpy30*-KO β-cells (Fig 2A) using calibrated chromatin immunoprecipitation sequencing (ChIP-seq) from FACS-enriched GFP-positive β-cells 45 days after tamoxifen administration. H3K4me1 also showed dramatic reduction (Fig 2A). Despite this, RNA-seq data showed a surprisingly modest reduction of *Ins1* and *Ins2* transcripts (Fig 2B). Therefore, robust expression of *Ins1* and *Ins2* RNA can occur in the absence of local H3K4me3. Notably, expression of several genes involved in insulin peptide maturation and packaging was downregulated (Fig S2A). For example *Nnat* and *Spcs3*, subunits of the signal peptidase complex which co-translationally converts preproinsulin to proinsulin ^34^, showed reduced expression in *Dpy30*-KO β-cells (Fig S2A-B). Proinsulin-to-insulin convertases *Pcsk1*, *Pcsk2*, and *Cpe* were not downregulated (Fig S2A). *Slc30a8*, encoding a zinc transporter required for maturation of insulin secretory granules ^35^ was highly downregulated in *Dpy30*-KO cells (Fig S2A). Insulin peptide content was reduced much more strongly than insulin RNA in *Dpy30*-KO islets (Fig 2B-C). Reduction of insulin content was confirmed in individual β-cells, where significant reduction to insulin granule size and density was seen in transmission electron micrographs of β-cells from *Dpy30*-KO mice 45 days after tamoxifen administration (Fig 2D-E). We conclude that loss of H3K4 methylation from mature β-cells impairs insulin biosynthesis.

**Figure 2.**
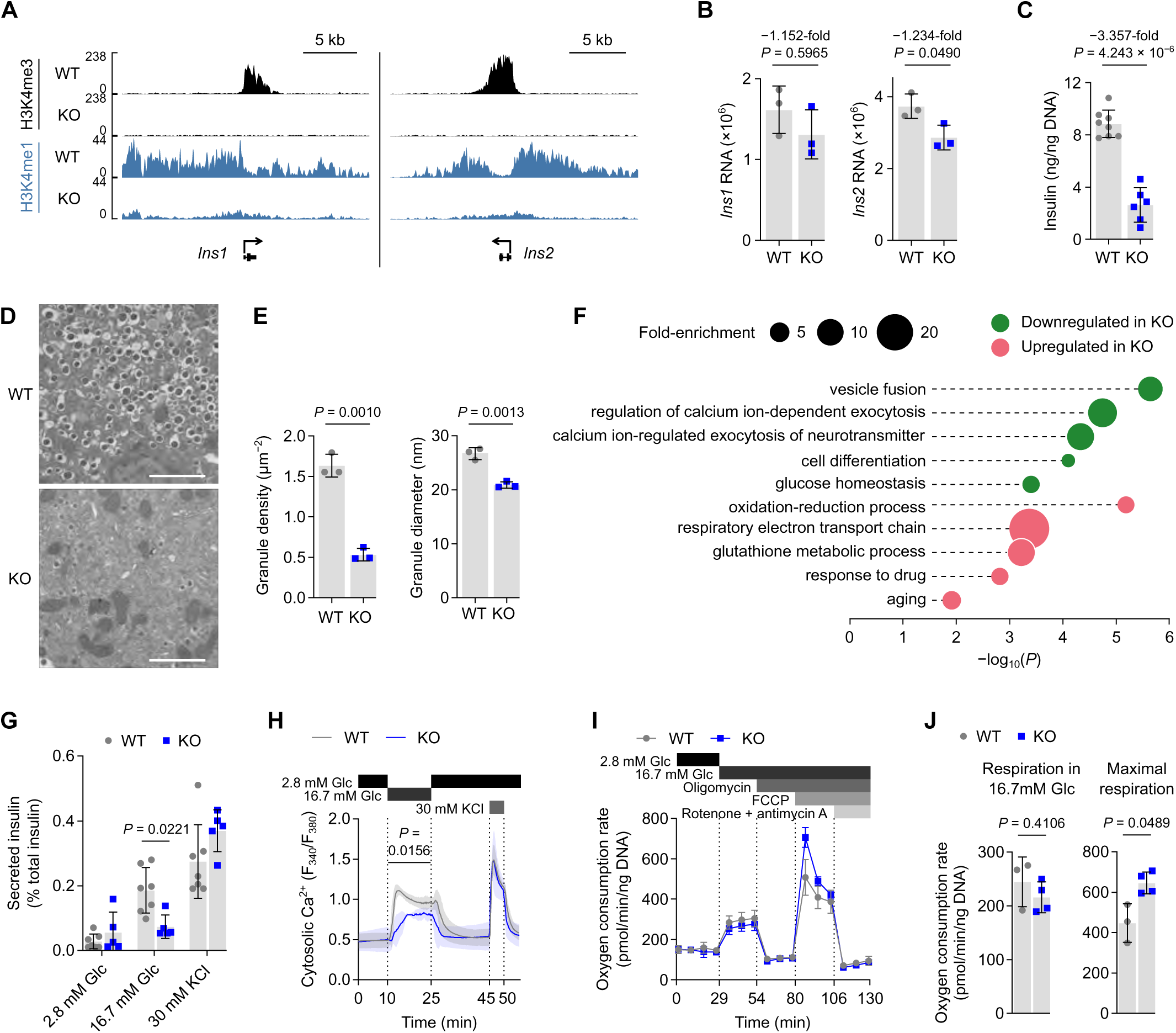
Genes involved in insulin production and glucose-induced activity are regulated by H3K4me3. **A,** Genome browser views of H3K4me3 and H3K4me1 enrichment at the *Ins1* and *Ins2* gene loci in *Dpy30*- WT and -KO β-cells 45-days after tamoxifen administration. **B,** *Ins1* and *Ins2* RNA levels in *Dpy30*-WT and -KO cells. Data presented as mean ± SD with values from individual mice (n = 3). Expression and *P*-values were calculated from RNA-Seq data using DESeq2 (Wald test with Benjamini-Hochberg correction). **C,** Insulin content in *Dpy30*-WT and -KO islets. Data presented as mean ± SD with values from individual mice (n = 8 WT, 6 KO). *P*-value calculated by two-tailed t-test with Welch’s correction. **D,** Representative transmission electron micrographs showing insulin granules in β-cells from a *Dpy30*-WT and a -KO mouse. Scale bars: 2 μm. **E,** Quantification of median insulin core granule size and density. Data presented as mean ± SD with values from individual mice (n = 3). *P*-values calculated by two-tailed t-test with Welch’s correction. **F,** Overrepresentation analysis of Gene Ontology: biological process terms in differentially expressed genes 45-days after tamoxifen administration. *P*-values represent EASE scores, a modified Fisher’s exact *P*-value, calculated using the Database for Annotation, Visualization, and Integrated Discovery (DAVID) v.6.8 ^87^. **G,** Insulin secretion from *Dpy30*-WT and -KO islets during static *in vitro* stimulation with glucose and KCl solutions, normalized to islet insulin content. *P*-values calculated by multiple two-tailed t-tests with Welch’s and Benjamini-Hochberg corrections; only values ≤ 0.05 are shown. **H,** Cytosolic Ca^2+^ concentration in islets from *Dpy30*-WT and -KO mice during *in vitro* perifusion of glucose and KCl solutions. *P*-values calculated for AUC in each time block by one-way ANOVA between genotypes (n = 4 WT, 3 KO); only values ≤ 0.05 are shown. **I,** Oxygen consumption rate in *Dpy30*-WT and - KO dispersed islet cells during treatment with the indicated compounds. n = 3 WT, 4 KO. **J,** Mitochondrial respiration in *Dpy30*-WT and -KO islet cells in 16.7 mM glucose and their maximal respiration capacity, inferred from the data shown in panel I. *P*-values calculated by two-tailed t-test with Welch’s correction (n = 3 WT, 4 KO).

Gene ontology analysis of genes showing reduced expression suggested the additional impairment of biological processes required for stimulated insulin secretion, namely vesicle fusion, calcium-dependent exocytosis, and glucose homeostasis (Fig 2F). Energy production processes including oxidation/reduction and the electron transport chain, meanwhile, were upregulated (Fig 2F). We therefore examined *ex vivo* islet activity during glucose stimulation. First, we measured insulin secretion as a fraction of insulin content to control for the 3.4-fold reduction on insulin in *Dpy30*-KO islets. Notably, the reduced insulin content of *Dpy30*-KO islets did not fully account for the impairment of insulin secretion. The persistent impairment was specific to high glucose stimulation, whereas secretion under nonstimulatory conditions, or bypassing glucose metabolism by depolarizing cells directly using KCl, showed no difference in insulin secretion between *Dpy30*-WT and -KO islets (after adjusting to total insulin content) (Fig 2G). Measurement of cytosolic calcium dynamics further supported an impairment of high glucose stimulated activity:. consistent with lower expression of calcium signaling genes (Fig 2F), high glucose-stimulated calcium influx showed a delayed response and lower magnitude in *Dpy30*-KO islets (Fig 2H). Cytosolic calcium concentration was not different in nonstimulatory conditions, and direct depolarization stimulated comparable calcium influx between islets from *Dpy30*-KO and *Dpy30*-WT mice (Fig 2H) further supporting a specific impairment in glucose responsiveness. We therefore wondered if *Dpy30*-KO impaired glucose catabolism. Expression of *Slc2a2*, encoding the glucose transporter GLUT2, was dramatically reduced in *Dpy30*-KO cells (Fig S2A-B), but expression of the rate-limiting enzyme of glycolysis, *Gck* ^36^, was not significantly altered (Fig S2A-B) nor was the rate of mitochondrial oxygen consumption in high glucose media altered (Fig 2I-J). In fact, mitochondrial respiratory capacity and area were increased in *Dpy30*-KO islets (Fig 2I-J, Fig S2C), consistent with elevated expression of oxidative phosphorylation genes in *Dpy30*-KO β-cells (Fig 2F). Therefore, glucose metabolism in β-cells was not appreciably impaired by reduction of H3K4 methylation. These results indicate that H3K4 methylation is essential for the expression of genes that couple glucose metabolism to insulin secretion in mature β- cells.

### Loss of H3K4me3 leads to gene downregulation in mature beta cells

It is notable that, while H3K4 methylation is well correlated with ^37^ and predictive of ^38^ transcription, our RNA-seq analysis indicates that globally only 4.3% of the 14,677 expressed genes were downregulated by depletion of H3K4 methylation in β-cells. This mirrors previous findings in yeast ^39^, *Drosophila* embryos ^15^, and mouse embryonic stem cells ^13, 14, 40^ that H3K4 methylation is not required for expression of most genes. We therefore wondered whether gene promoters that are sensitive to H3K4 methylation in β-cells have generalizable features. We first confirmed that H3K4me3 and H3K4me1 were depleted at active gene promoters in *Dpy30*-KO chromatin (Fig 3A). We further confirmed genome-wide reduction of H3K4me3 and H3K4me1 by comparing enrichment between WT and *Dpy30*-KO chromatin in 10-kb bins spanning the genome (Fig 3B-C), confirming that *Dpy30* is necessary for maintenance of β-cell H3K4 methylation. Despite this, the transcriptomes of *Dpy30*-KO and -WT cells were well correlated, and the total RNA expression of *Dpy30*-KO cells was not decreased (Fig 3D). Therefore, global depletion of H3K4 methylation does not cause a global decrease in gene expression in mature β-cells.

**Figure 3.**
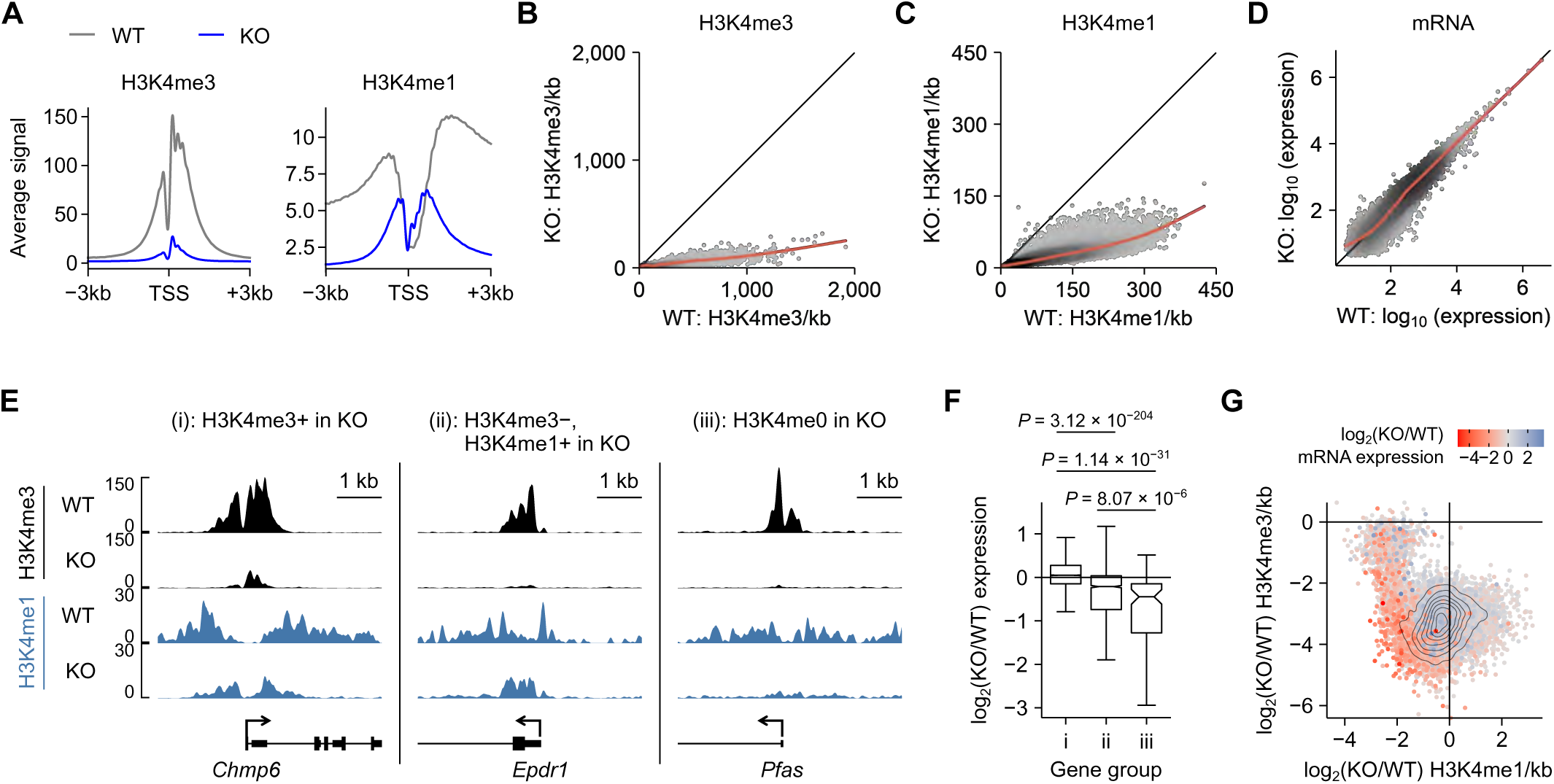
TSS-associated H3K4 methylation maintains gene expression in mature beta cells. **A,** Average enrichment profiles of H3K4me3 and H3K4me1 at the TSS of all expressed genes in *Dpy30*-WT and -KO chromatin. **B-C,** Scatterplot of H3K4me3 (B) and H3K4me1 (C) enrichment in 10-kb bins spanning the genome in *Dpy30*-KO versus -WT cells. **D,** Scatterplot showing spike-in normalized gene expression in *Dpy30*-KO versus -WT cells. **E,** Genome browser views show examples of genes that retain H3K4me3 (i), lose H3K4me3 but retain H3K4me1 (ii), or lose H3K4me3 and H3K4me1 (iii) from the TSS in *Dpy30*-KO chromatin. **F,** Box and whisker plot showing log2(fold-change) of RNA expression for gene groups described in panel E. *P* indicates the Wilcoxon signed-rank test result with Benjamini-Hochberg correction. **G,** Contour scatterplot showing log2(fold-change) of H3K4me1 enrichment (x-axis) plotted against log2(fold-change) of H3K4me3 enrichment (y-axis) in the TSS ±1 kb of all expressed genes. The log2(fold- change) of RNA expression of the associated genes is shown as a colour gradient.

To explore further, we segregated genes into three groups based on residual enrichment for H3K4 methylation, determined using MACS2 ^41^. Group (i) retained at least some H3K4me3 enrichment at the transcription start site (TSS) (10,045 genes), group (ii) lost H3K4me3 but retained at least some H3K4me1 enrichment (2,302 genes), and group (iii) lost H3K4me3 and H3K4me1 (111 genes) (Fig 3E). As a group, genes that retained at least some enrichment for H3K4me3 at the TSS were not generally downregulated in *Dpy30*-KO cells (Fig 3F). Loss of both H3K4me3 and H3K4me1 from the TSS was linked to significant impairment of gene expression, and genes that lost H3K4me3 but retained H3K4me1 showed an intermediate phenotype (Fig 3F). To visualize this relationship in a threshold-free manner, we plotted the change in H3K4me3, H3K4me1, and RNA expression for all active TSSs (Fig 3G). The majority of genes clustered in the bottom left quadrant, indicating some reduction of both H3K4me3 and H3K4me1 from the TSS. Strikingly, genes that were highly downregulated (red) clustered at the extreme edge of the distribution, indicating that genes experiencing the greatest relative loss of both H3K4me3 and H3K4me1 were highly downregulated (Fig 3G). Together, these data support a role for H3K4me3 in contributing to active gene expression: while the genome-wide reduction of H3K4me3 did not influence total cellular mRNA content, excess loss of H3K4me3 on a per-gene basis leads to downregulation of that gene.

### H3K4me3 enforces expression of weakly active and repressed genes in mature beta cells

We then compared the chromatin profile of downregulated and stably expressed genes by measuring H3K27me3, a marker of developmentally repressed chromatin, and H3K27ac, a marker of active chromatin, using ChIP-seq. The downregulated gene set had a weakly active profile even in WT cells: enrichment for H3K4me3 and H3K27ac were low and H3K4me1 and H3K27me3 were high in comparison to stably expressed genes (Fig 4A-B). Accordingly, the downregulated gene set showed lower absolute expression in WT cells than stably expressed genes (Fig 4C). The DNA sequence of downregulated genes also tended to be relatively GC rich (Fig 4E). Imprinted genes, comprising genomic loci for which one allele is silenced by DNA methylation, tended to be downregulated in *Dpy30*-KO cells (Fig 4F), raising the possibility that H3K4me3 impedes *de novo* DNA methylation at some genes in mature β-cells, as it does in mouse embryonic stell cells ^42^. To summarize, genes that lose an especially large fraction of H3K4 methylation tend to be downregulated in *Dpy30*-KO cells; they tend to already be in a weakly active state in WT β-cells and may be more susceptible to silencing in the absence of H3K4 methylation.

**Figure 4.**
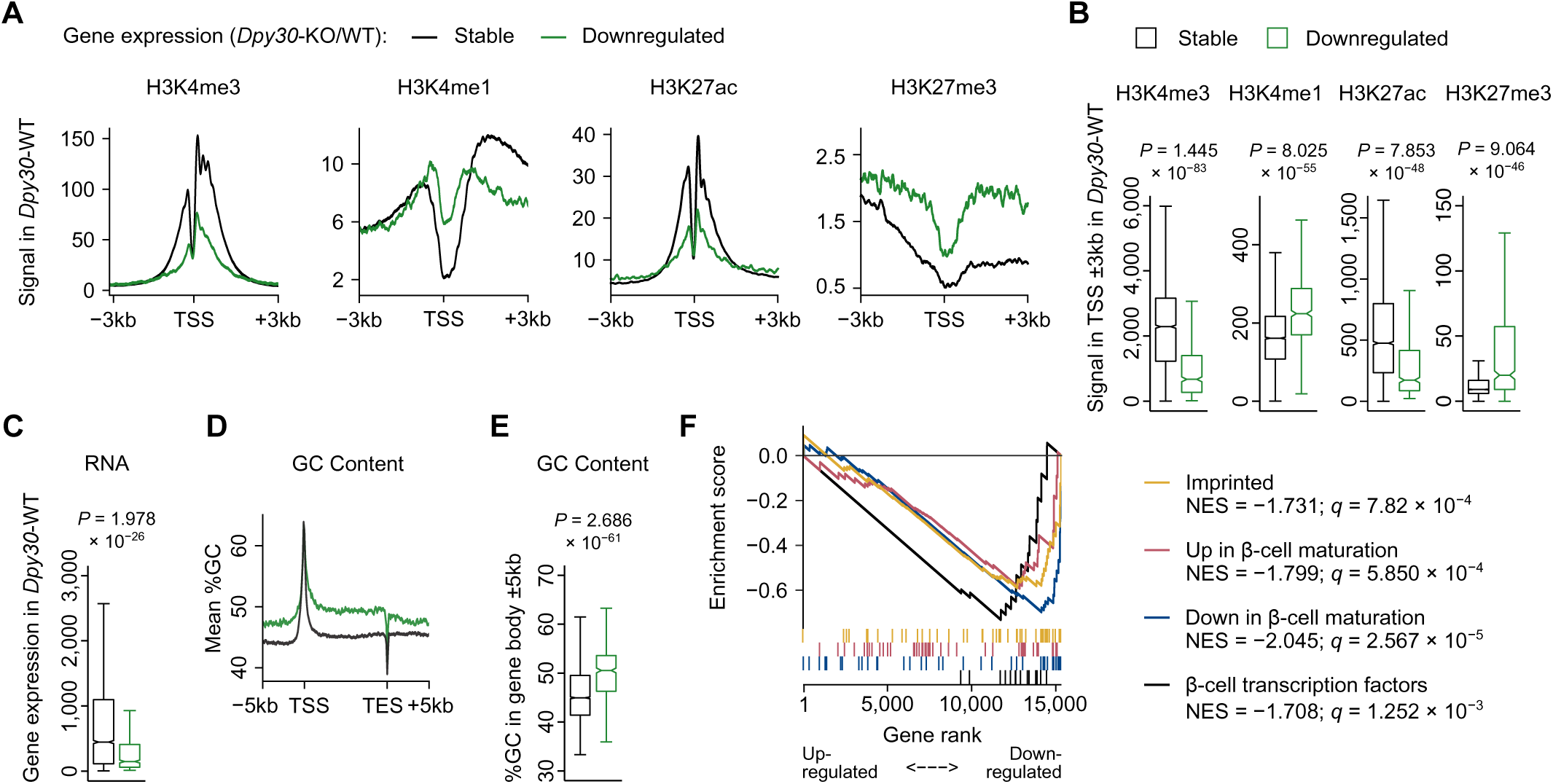
Weakly active and developmentally regulated genes are susceptible to downregulation in H3K4me3-deficient beta cells. **A,** Average enrichment profiles of H3K4me3, H3K4me1, H3K27ac, and H3K27me3 in *Dpy30*-WT cells at TSSs of genes that are stably expressed or downregulated by *Dpy30*-KO. **B,** Box and whiskers plots showing quantification of data shown in panel A. **C,** Box and whiskers plot showing mRNA expression in *Dpy30*-WT cells for genes stably expressed or downregulated by *Dpy30*-KO. **D,** Average %GC content profile in the gene body ±5 kb of genes that are stably expressed or downregulated by *Dpy30*-KO. **E,** Box and whiskers plot showing quantification of data shown in panel D. **F,** Running enrichment plot for imprinted genes (ref.^88^) and genes that are up- or downregulated during mouse β-cell maturation (adults versus postnatal day 10, ref. ^46^), and mature β-cell transcription factor genes. NES: normalized enrichment score. *P* values calculated using Wilcoxon signed-rank test (panels B,C,E) or permutation test (F) with Benjamini-Hochberg correction.

### H3K4me3 deficiency leads to a less active and more repressed epigenome in mature beta cells

Our finding that H3K4 methylation is especially important for maintaining expression of weakly expressed genes is intriguing in light of a recent report by Dobrinić et al. (ref ^43^) that demonstrated Polycomb-mediated gene repression especially constrains weakly expressed genes ^43^. The trithorax group was originally defined as antagonistic to Polycomb-mediated repressive activity ^44^. During development, repressive (H3K27me3) and active (H3K4me3) states are dynamically regulated and resolution of the genome into active and repressive domains is fundamental to the establishment of terminal cell-type- specific transcriptomes ^45^. As such, it is notable that expression of genes that are dynamically regulated during, and important for, β-cell maturation ^46^ tend to be lower in *Dpy30*-KO cells (Fig 4F). This led us to ask whether H3K27me3 expands in H3K4me3-deficient β-cells. As shown in Fig 5A, the large-scale distribution of H3K27me3-positive compartments is unchanged in *Dpy30*-KO chromatin. H3K7me3 enrichment in 10-kb windows across the genome remained similar between *Dpy30*-KO and -WT (Fig 5B), together indicating that global reduction of H3K4me3 and H3K4me1 did not result in expansion of Polycomb-repressed compartments. However, H3K27me3 enrichment specifically increased at promoters of genes that were downregulated in *Dpy30*-KO cells (Fig 5C-E). These data indicate that genome-wide H3K27me3 organization is largely insensitive to reduction of H3K4 methylation in this model; however, maintenance of H3K4 methylation is necessary to prevent targeted Polycomb gene repression in mature β-cells.

**Figure 5.**
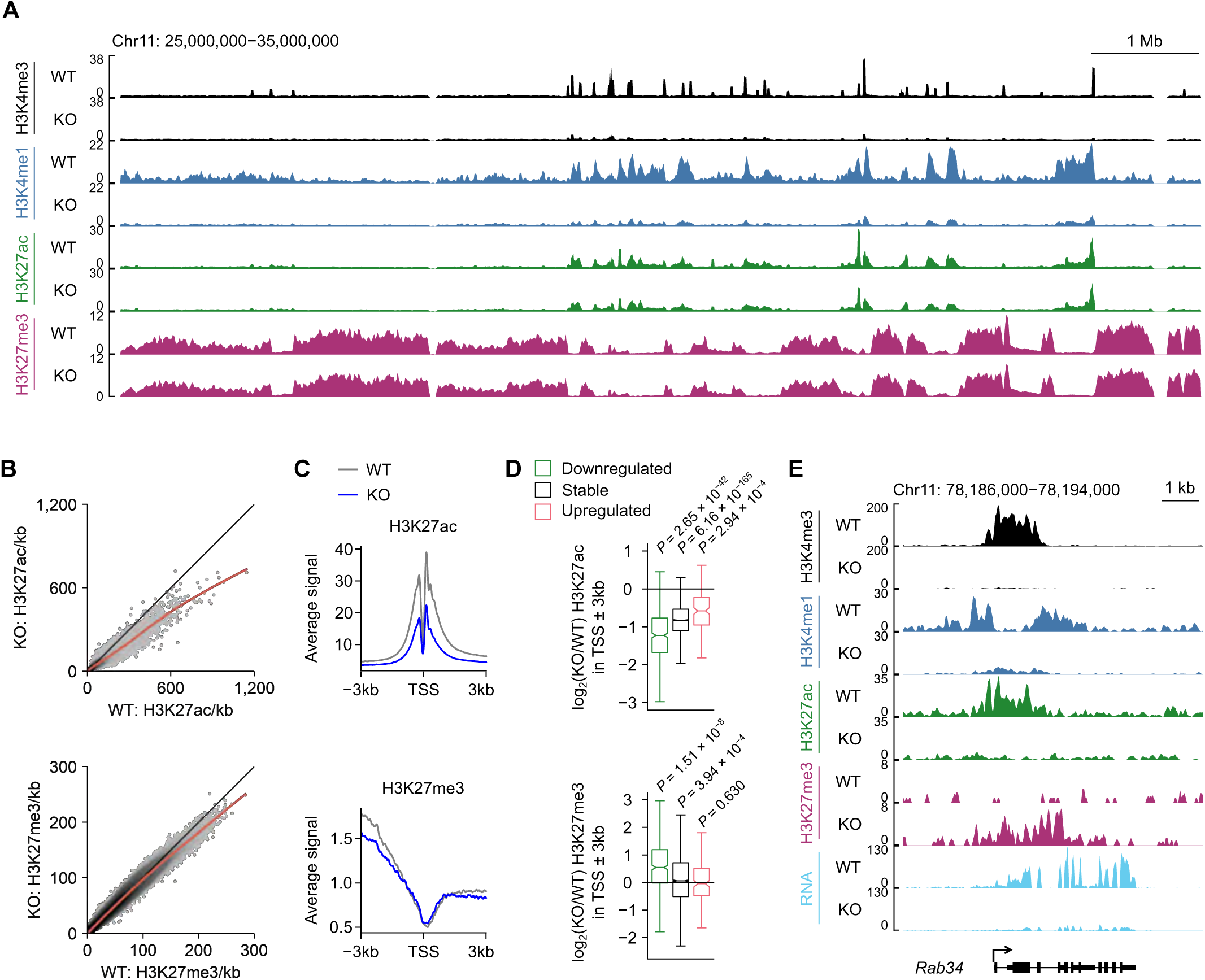
H3K4me3 deficiency leads to a less active and more repressed epigenome in mature beta cells. **A,** Genome browser view of H3K4me3, H3K4me1, H3K27ac, and H3K27me3 in *Dpy30*-WT and -KO cells showing preservation of mega base-scale organization of H3K27me3-positive PcG compartments. **B,** Scatterplots of H3K27ac (top) and H3K27me3 (bottom) enrichment in 10-kb bins spanning the genome in *Dpy30*-KO versus -WT cells. **C,** Average enrichment profiles of H3K27ac (top) and H3K27me3 (bottom) at the TSS of all expressed genes in *Dpy30*-WT and -KO cells. **D,** Box and whisker plots showing the log2(fold- change) of H3K27ac (top) and H3K27me3 (bottom) enrichment in the TSS of transcriptionally downregulated, stable, or upregulated genes in *Dpy30*-KO compared to -WT cells. *P*-values calculated with Wilcoxon rank sum test with Benjamini-Hochberg correction. **E,** Genome browser view of a downregulated gene *Rab34* showing loss of H3K4me3, H3K4me1, H3K27ac, and RNA, and accumulation of H3K27me3, in *Dpy30*-KO cells.

In contrast to the locus-specific accumulation observed of H3K27me3, H3K27ac, a marker of active chromatin, was diminished throughout the genome in *Dpy30*-KO cells (Fig 5B). Like H3K4me3, H3K27ac was reduced even at promoters of genes that were stably expressed or upregulated in *Dpy30*- KO cells (Fig 5D), indicating that the generalized decrease in H3K27ac was not a consequence of gene downregulation. Despite the global reduction, however, H3K27ac was still positively associated with gene expression in *Dpy30*-KO chromatin: H3K27ac peaks gaining intensity were near upregulated genes, and H3K27ac peaks losing intensity were near downregulated genes, more frequently than expected by chance (Fig S3). These observations are consistent with a model wherein acetylation of H3K27 is partially a consequence of H3K4 methylation ^14^ and of transcription ^47^.

### Reduction to transcriptional consistency in *Dpy30*-KO beta cells

H3K4me3 enrichment has also been linked with transcriptional consistency—that is, low variability, or “noise”, in the expression of a given gene between cells ^48, 49^. A related metric, transcriptional entropy, is a measure of the “specialization” of a cell’s transcriptome, where a low entropy describes a narrow distribution of highly expressed genes in each cell, versus a high entropy where gene expression becomes less predictable ^50^. In islets, transcriptional entropy decreases during maturation ^50^ and increases in T2D ^4^. As such, we asked whether reduction of H3K4 methylation impairs transcriptional consistency in mature β-cells. To measure transcriptional consistency, we processed islets from a WT and *Dpy30*-KO mouse for single cell RNA-seq 45-days after tamoxifen administration, generating 2,510 WT and 2,597 KO cell transcriptomes that passed quality filtering and that were identified as β-cells based on unsupervised clustering and high expression of *Ins1 and Ins2* (Fig S4A). A population of β-cells expressing *tdTomato*, indicating that they were not Cre-recombined, formed a distinct cluster from recombined *GFP*-positive β- cells from the *Dpy30*-KO mouse (Fig 6A). To model divergence of *GFP*-positive from *tdTomato*-positive β- cells (hereafter, mG and mT cells, respectively), we performed pseudotime ordering using the mT cluster as the ground state (Fig 6B). As seen in Fig 6C, *Ins1* and *Ins2* expression became notably more stochastic over pseudotime. To quantify variability of global transcription, we generated transcriptomic entropy scores for each β-cell. In *Dpy30*-KO β-cells, there is an overall increase in transcriptional entropy along pseudotime (Fig 6D) and in mG cells compared to mT cells (Fig 6E). This supports a role for H3K4 methylation in maintaining high transcriptional consistency in mature β-cells. Since higher entropy is a feature of immature β-cells ^50^, we examined expression of genes associated with immaturity and dedifferentiation and again did not observe any induction of these genes in Dpy30-KO cells (*c.f.* Figs 4F & S4B-C). Therefore, *Dpy30*-KO in mature β-cells led to a more stochastic gene expression profile but did not alter cell fate or re-activate an immature cell state.

**Figure 6.**
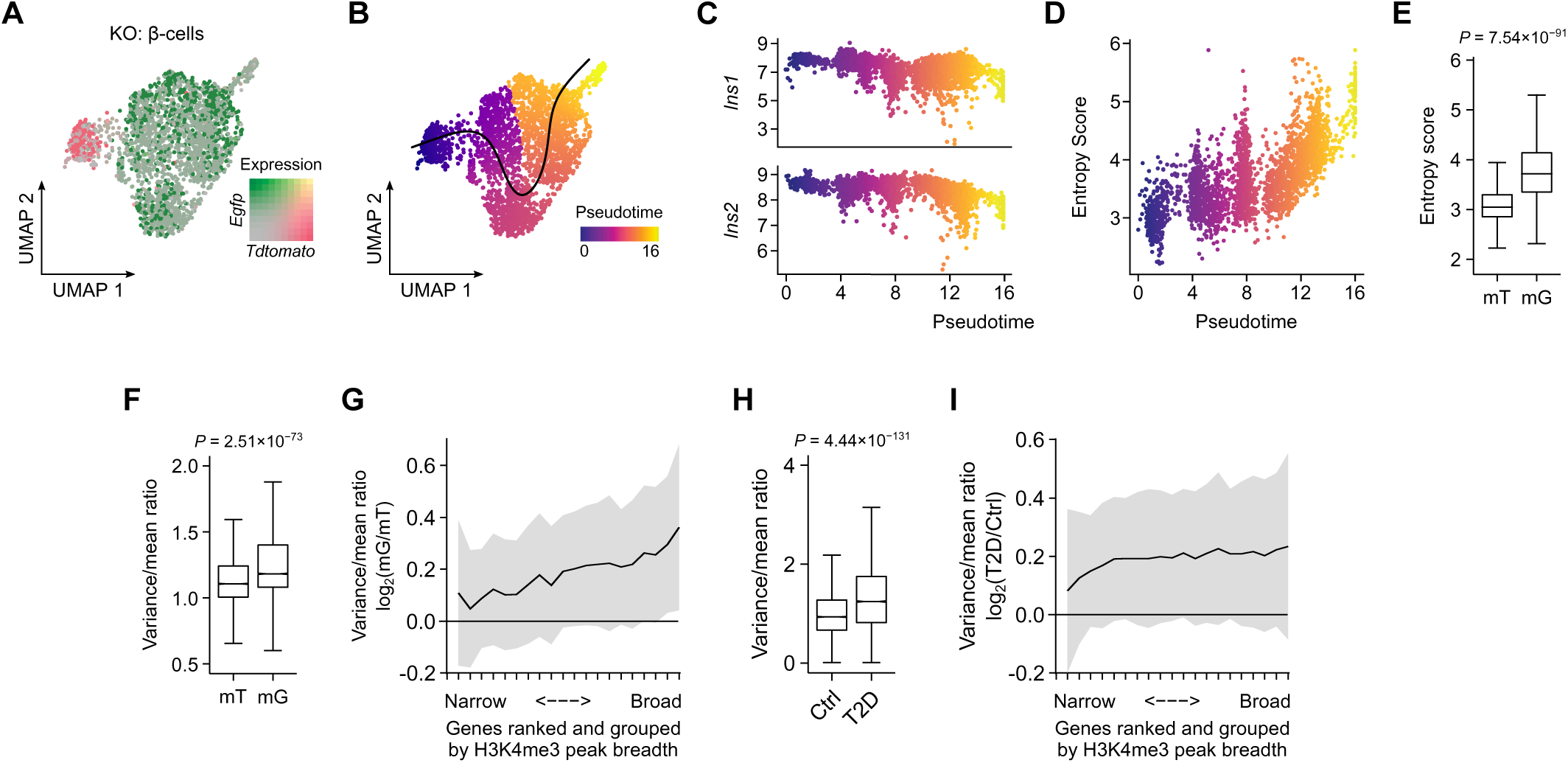
Reduction of transcriptional consistency in *Dpy30*-KO and diabetic beta cells. **A,** UMAP visualization of β-cell transcriptomes from a *Dpy30*-KO mouse. Expression of *Tdtomato*, *Egfp*, and their overlap are shown as a colour gradient. **B,** Pseudotime representation of A. **C,** Scatterplot showing *Ins1* and *Ins2* expression in β-cells plotted against pseudotime. **D,** Scatterplot showing gene expression entropy scores of β-cells plotted against pseudotime. **E,** Box and whisker plot showing the transcriptional entropy of β-cells in *Tdtomato+* and *Egfp+* clusters (n = 274 and 2,323 cells, respectively). **F,** Box and whisker plot showing the variance/mean ratio of gene expression across β-cells in *Tdtomato+* and *Egfp+* clusters. **G,** Variance/mean ratio of β-cell gene expression in *Egfp+* versus *Tdtomato+* clusters (y-axis) plotted as a function of H3K4me3 breadth in quantiles (x-axis) (622 genes/quantile). Data presented as mean ± SD. **H-I,** same as F-G in β-cells from human donors with or without T2D, using H3K4me3 ChIP-seq data from GSE50386 (ref. ^51^) and scRNA-seq data from GSE124742 ref. ^52^).

We then asked whether the loss of transcriptional consistency was linked to the initial breadth of the H3K4me3 peak, since it has been suggested that H3K4me3 peaks that span an especially large stretch of DNA (i.e. broad peaks) may enforce high transcriptional consistency of associated genes ^48, 49^. We hypothesized that genes with broader H3K4me3 peaks are more susceptible to increasingly stochastic expression when H3K4me3 levels are reduced, compared to other genes with narrower peaks. To test this, we ranked H3K4me3 peaks in WT cells in order of increasing breadth, grouped them into 20 quantiles, each representing 5% (622) of H3K4me3 peaks with an associated gene. We then compared the variance in expression, normalized to absolute expression level, of genes in each quantile between mT and mG cells. Overall transcriptional variability between cells was increased in mG cells (Fig 6F). There was a positive relationship between H3K4me3 peak breadth and the change in transcriptional variability, where greater H3K4me3 peak breadth was linked to greater gain of variability in *Dpy30*-KO cells (Fig 6G). This supports a role for H3K4me3 in constraining transcriptional variability in β-cells, which is positively linked to peak breadth. Intriguingly, a similar pattern emerges in β-cells from human donors with T2D. Analysis of publicly available datasets for H3K4me3 ChIP-seq from healthy human β-cells (GSE50386, ref. ^51^) and scRNA-seq from human donors with or without T2D (GSE124742, ref. ^52^) showed that variability of gene expression was higher in β-cells from donors with T2D (Fig 5H) and that the gain in variability is positively correlated with initial H3K4me3 peak breadth (Fig 5I).

### H3K4me3 peak breadth stratifies genes dysregulated in type 2 diabetes

We further explored the relationship between H3K4me3 peak breadth, gene expression, and diabetes. H3K4me3 forms 18,936 distinct peaks with a median breadth of 1.2 kb in WT mouse β-cells. Ranking H3K4me3 peaks by breadth revealed a class of exceptionally broad peaks spanning up to 21 kb, including those at essential β-cell-enriched transcription factor genes (Fig 7A), consistent with the idea that genes important for maintaining cell identity are marked with broad H3K4me3 peaks ^48, 53^. As expected for repressed genes, the majority (29 of 39, or 74%) of islet “disallowed genes” ^54^, meanwhile, are H3K4me3- negative in healthy β-cells. Gene expression was moderately correlated with promoter H3K4me3 peak breadth (Spearman rank correlation R = 0.4287); however, genes with exceptionally broad peaks did not have exceptionally high RNA expression (Fig S5A). For example, Fig 7B shows the H3K4me3 profiles of a housekeeping gene with a typical H3K4me3 peak (*Rplp0*), and an expression-matched β-cell-enriched transcription factor with a broad peak (*Nkx6-1*). Therefore, β-cell lineage-enriched genes are marked by broad H3K4me3 peaks.

**Figure 7.**
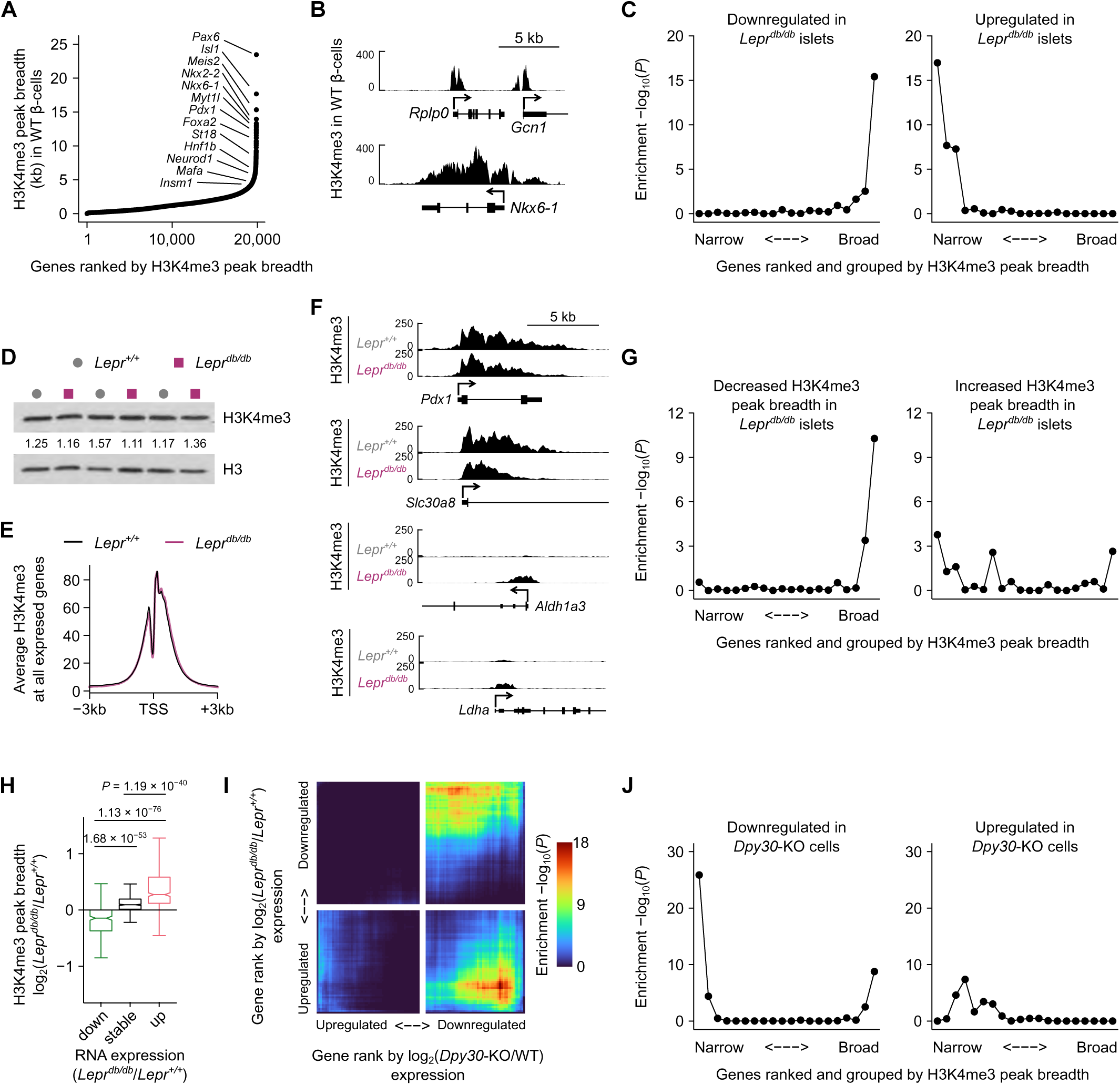
H3K4me3 peak breadth encodes gene expression dynamics in a mouse model of type 2 diabetes. **A,** H3K4me3 peaks in *Dpy30*-WT β-cells ranked from narrow to broad. Peaks associated with β-cell transcription factor genes are labeled. **B,** Genome browser views showing H3K4me3 enrichment in WT β- cells at a housekeeping gene (*Rplp0*), and an expression-matched β-cell transcription factor gene (*Nkx6-1*). **C,** Enrichment *P*-values (y-axis) of genes down- or upregulated in islets from *Lepr^db/db^* mice, where genes have been ranked and grouped by H3K4me3 peak breadth from narrow to broad (x-axis). Each group contains 622 genes. *P*-values were calculated using one-sided Fisher’s exact test. **D,** Immunoblots of H3K4me3 and total histone H3 in islet lysate from *Lepr^+/+^* and *Lepr^db/db^* mice. H3-normalized H3K4me3 band densities are listed. **E,** Average TSS enrichment profile of H3K4me3 in *Lepr^+/+^* and *Lepr^db/db^* islets. **F,** Genome browser views of H3K4me3 in *Lepr^+/+^* or *Lepr^db/db^* islets at notable genes that are downregulated (*Pdx1*, *Slc30a8*) or induced (*Aldh1a3*, *Ldha*) in *Lepr^db/db^* islets. **G,** Enrichment *P*-values of H3K4me3 peaks showing significant reduction or expansion of peak breadth in *Lepr^db/db^* versus *Lepr^+/+^* islets (y-axis) in the different gene groups ranked from narrow to broad (x-axis). Enrichment *P*-values were calculated using one-sided Fisher’s exact test. **H,** Box and whisker plots showing the log2(fold-change) of H3K4me3 peak breadth for genes that are downregulated, stably expressed, or upregulated in *Lepr^db/db^* islets. *P*-values calculated with Wilcoxon rank sum test with Benjamini-Hochberg correction. **I,** Stratified rank/rank hypergeometric overlap plot comparing gene expression changes caused by *Dpy30*-KO (x-axis) and by *Lepr^db/db^* (y-axis) mutations. Colourscale shows the enrichment *P-*value calculated using the RRHO2 R package ^59^**. J,** Same as panel C for genes that are differentially expressed in *Dpy30*-KO cells.

To explore the relationship between H3K4me3 peak breadth and transcriptome remodeling associated with diabetes, we turned to a mouse model of T2D. Leptin receptor-deficient *Lepr^db/db^* mice exhibit overfeeding, obesity, fasting hyperglycemia, and glucose intolerance, along with β-cell dysfunction^55^. We retrieved a published bulk RNA-seq dataset from *Lepr^db/db^* and *Lepr^+/+^* islets ^56^ and compared H3K4me3 peak breadth with gene expression changes. Interestingly, genes that become downregulated in *Lepr^db/db^* islets were explicitly enriched in the group of genes with broad H3K4me3 peaks (Fig 7C, left). Furthermore, housekeeping genes (GSEA:M11197, ref ^57^), genes involved in endocrine pancreas development (GSEA:M12875), and maturity onset diabetes of the young (MODY) genes (GSEA:M18312) were enriched in the broadest H3K4me3 peaks (Fig S5B). In contrast, genes that are upregulated in *Lepr^db/db^* islets were uniquely enriched in the narrowest H3K4me3 peaks (Fig 7C, right). As H3K4me3 peak breadth was positively correlated with expression level (Fig S5A), we tested whether these observations could be secondary to a relationship wherein highly expressed genes become downregulated, and weakly expressed genes become upregulated, in *Lepr^db/db^*. Ranking genes by RNA expression level instead of H3K4me3 peak breadth proved less effective at stratifying each gene set except housekeeping genes (Fig S5B), supporting a primary link with H3K4me3. Therefore, genes that are differentially expressed in *Lepr^db/db^* islets are diametrically stratified according to H3K4me3 breadth, with upregulated genes having narrow peaks and downregulated genes having broad peaks (in non-diabetic conditions). These observations are replicable in humans using publicly available H3K4me3 ChIP-seq data of β-cells from donors without diabetes (GSE50386, ref ^51^) and RNA-seq data of islets from donors with and without T2D (GSE50398, ref ^58^) (Fig S5C-E).

### H3K4me3 peak breadth dynamics encode gene expression changes in a mouse model of type 2 diabetes

The positive association between H3K4me3 peak breadth and differential gene expression in T2D prompted us to ask if H3K4me3 is altered in diabetic islets. Immunoblots showed that, overall, H3K4me3 levels were not altered in islets from *Lepr^db/db^* mice (Fig 7D). Calibrated ChIP-seq similarly showed that average enrichment for H3K4me3 across all active promoters is remarkably similar between *Lepr^+/+^* and *Lepr^db/db^* islets (Fig 7E). While we did not find evidence for a global shift in H3K4me3 levels, we observed contraction of H3K4me3 at downregulated genes, for example the transcription factor *Pdx1* and zinc transporter *Slc30a8*, and expansion of H3K4me3 at upregulated genes, for example the disallowed *Aldh1a3* and *Ldha* (Fig 7F, Fig S5F). In general, broad and narrow H3K4me3 peaks showed significant tendency to shrink or expand, respectively (Fig 7G). Similarly, genes that were transcriptionally up- or downregulated in *Lepr^db/db^* islets showed corresponding expansion or contraction of HK4me3 levels, respectively (Fig 7H). Therefore, a pattern emerges in *Lepr^db/db^* islets wherein weakly active genes tend to gain H3K4me3 and become upregulated, while genes with broad H3K4me3 peaks tend to lose some H3K4me3 and become downregulated.

Finally, we asked whether genes that are dysregulated in *Lepr^db/db^* islets are regulated by H3K4 methylation, i.e., are downregulated in the day-45 *Dpy30*-KO RNA-seq. To test the overlap between differential gene expression in the *Dpy30*-KO and *Lepr^db/db^* models, we used a threshold-free rank-rank hypergeometric overlap test (RRHO) ^59^. There was significant overlap between genes downregulated in *Dpy30*-KO mice with genes that were differentially expressed (either up or down) in *Lepr^db/db^* mice (Fig 7J). Thus, H3K4 methylation helps maintain expression of genes that become dysregulated in the diabetic *Lepr^db/db^* model. Accordingly, genes downregulated in *Dpy30*-KO mice were enriched in both the narrow and broad H3K4me3 peak groups (Fig 7I-J), thus mirroring the pattern for dysregulated genes in the *Lepr^db/db^* model (Fig 7C). This observation in *Dpy30*-KO cells is in agreement with observations presented above: that weakly active genes are most downregulated in *Dpy30*-KO (Fig 4), and that β-cell transcription factor genes have broad peaks (Fig 7A) and tend to be downregulated by *Dpy30*-KO (Fig 4F). Together, this supports the idea that differential gene expression between *Lepr^+/+^* and *Lepr^db/db^* mice is partially driven by changes in H3K4me3 peak breadth.

## Discussion

Here we set out to address whether H3K4me3 is required for maintenance of gene expression in mature β-cells. To this end we deleted a core component of the COMPASS H3K4 methyltransferase complexes, *Dpy30*, leading to dramatic genome-wide reduction of H3K4me3 in mouse β-cells. Loss of H3K4 methylation led to a less active and more repressed epigenome profile which locally correlated with gene downregulation. Gene expression became more stochastic, a feature also shared with T2D. H3K4me3- dependent gene expression meaningfully contributed to maintenance of mature β-cell function and maturity. H3K4me3-deficient cells showed loss of maturity including reduced glucose stimulated insulin secretion, reduced expression of terminal markers, and higher transcriptome entropy. These analyses support a role for H3K4me3 in maintaining β-cell gene expression and function.

In *Dpy30*-KO cells, changes to RNA expression were linked to local changes to histone methylation, leading to significant dysregulation of around 5% of genes, but overall cellular RNA levels were not reduced. H3K4 methylation may, therefore, be important for allocation of transcriptional machinery to particular genes, rather than their absolute activity. In other words, promoter-associated H3K4me levels modulate the expression of target genes but global H3K4me3 or H3K4me1 levels should not be taken as a reflection of a cell’s transcriptional output. We note, however, that total mature RNA content, which we measured here, is not a direct measure of transcriptional activity, and measurement of nascent RNA may have uncovered more widespread roles in transcription regulation.

Hinting at more general roles in transcription is the observation that *Dpy30*-KO β-cells displayed a more stochastic gene expression profile. High gene expression variability between cells in a tissue is a beneficial feature in some circumstances – it can increase adaptability to stress and can reflect a high degree of cell subtype specialization. In other circumstances, high variability is associated with aging and disease ^60–63^, and we find that it is increased in β-cells from donors with T2D. Very broad H3K4me3 peaks are proposed to confer high transcriptional stability ^48^, and experimental perturbation of H3K4me3 using *Dpy30*-KO supports this idea. Further, transcription of genes with H3K4me3 peaks of any breadth tended to become more variable in *Dpy30*-KO, meaning that H3K4me3 also generally confers consistent mRNA transcription. Defining exactly how H3K4me3 influences transcriptional activity of select genes will require detailed consideration of the larger epigenetic environment and RNA polymerase activity.

Since H3K4 cannot be simultaneously mono- and trimethylated, enrichment of H3K4me3 causes H3K4me1 to be generally excluded near TSSs. The degree of exclusion relates to the transcriptional output of the TSS, where weakly expressed genes display closer infiltration of H3K4me1 ^64^. In *Dpy30*-KO cells, replacement of H3K4me3 with H3K4me1 in the TSS was associated with transcriptional downregulation, consistent with a report that promoter-associated H3K4me1 is repressive ^65^. Surprisingly, replacement of H3K4me3 with H3K4me1 buffered gene downregulation compared to complete demethylation, suggesting that promoters with H3K4me1 are not repressed but simply less active, and H3K4me1 can partially compensates for loss H3K4me3 in promoters. This mirrors a finding that enhancers can function with H3K4me3 instead of H3K4me1 ^13^. These results argue against an immutable functional distinction between H3K4me1 and H3K4me3, at least in this model.

One of the most surprising findings is that promoters in a weakly active state, with little H3K4me3, show the greatest reliance on H3K4 methylation for the maintenance of gene expression. It. A possible explanation is that these promoters are particularly targeted for epigenetic silencing ^66^ since H3K4 methylation sterically hinders repressive H3K27 and DNA methyltransferase activity ^42, 67^. Genes that were downregulated in *Dpy30*-KO cells showed high initial enrichment of H3K27me3 and further accumulation after loss of H3K4me3. Weakly-active genes are also the most likely to be upregulated after inactivation of the Polycomb system ^43^ and in β-cells, inactivation of the Polycomb system leads to activation of previously silenced genes ^4^. Therefore, weakly active and suppressed genes may be the most responsive targets in ongoing competition between Polycomb and trithorax systems in mature β-cells. Future studies may unravel the degree to which H3K4me3 is an activating versus anti-repressive signal *in vivo*.

Peak breadth has emerged as an informative dimension of H3K4me3 enrichment: H3K4me3 peak breadth predicts gene expression level ^38^; breadth dynamics predict gene expression dynamics during methionine restriction *in vivo* and *in vitro* ^68^; and, exceptionally broad H3K4me3 peaks in a given cell type are predictive of genes critical to the maintenance of that cell’s lineage ^48^. From our analyses of peak breadth, we confirm that terminal markers of β-cells, including *Ins1/2* and mature β-cell enriched transcription factor genes, are marked with exceptionally broad H3K4me3 peaks in mature β-cells. *Dpy30*- KO led to nonspecific reduction of H3K4me3 genome-wide but transcriptional downregulation was largely restricted to genes that originally had either very narrow, or very broad, H3K4me3 peaks, indicating that these two classes of promoters in particular rely on H3K4me3 for the maintenance of gene expression. Compared to the effect at narrow peaks described above, the transcriptional defects for genes with broad H3K4me3 enrichment was more modest. For example, *Ins1* and *Ins2* were downregulated by 15 to 25%, remaining the two most highly expressed genes in *Dpy30*-KO cells despite suffering complete loss of their broad H3K4me3 peaks. Since broad peaks colocalize with many other activating chromatin marks ^48, 53^, the modest effect likely results from compensation from other regulatory inputs. In any case, our results show that broad H3K4me3 increases expression of associated genes but is not strictly required for their transcription.

The *Dpy30*-KO model highlighted the central importance of relative H3K4me3 levels on a per-gene basis rather than absolute enrichment; thus, reduction of peak breadth may exert a stronger impairment on gene expression in scenarios where H3K4me3 is not simultaneously reduced at all other genes. To that point, we find that promoters with broad H3K4me3 peaks showed a unique tendency to shrink, and for the associated gene to be downregulated, in *Lepr^db/db^* islets. Meanwhile, promoters marked with narrow H3K4me3 peaks showed a significant tendency to be upregulated and gain H3K4me3 in *Lepr^db/db^* islets. It is notable that these two classes of promoters are the same that are most sensitive to H3K4me3 enrichment. More work will be required to determine what drives H3K4me3 peak dynamics in *Lepr^db/db^* islets and to what extent those mechanisms are active in human T2D. In any case, these results favour a model wherein redistribution of H3K4me3 away from promoters of genes with broad H3K4me3 peaks, that enforce β-cell identity or functions, in favour of relatively inactive, weakly-expressed genes, including developmentally repressed and disallowed genes, contributes to transcriptome remodeling in diabetes.

## Methods

### Animal strains and care

Mice harboring conditional alleles of *Dpy30* (derived from EUCOMM EM:09575 ^21^) and *Rosa26^mTmG^* (Jax 007576 ^29^) were crossed with *Pdx1-CreER^Tg^* (Jax 024968 ^28^) or *Ins1^Cre^* (Jax 026801 ^33^) lines. Genotypes used for *Pdx1-CreER* studies were: KO: *Pdx1-CreER^Tg/^*^0^; *Dpy30^flox/flox^*; *Rosa26^mTmG/+^*; WT: *Pdx1-CreER^Tg/^*^0^; *Dpy30^+/+^*; *Rosa26^mTmG/+^* or *Pdx1-CreER*^0^*^/^*^0^; *Dpy30^flox/flox^*; *Rosa26^mTmG/+^*. Genotypes used for *Ins1^Cre^* studies were: KO: *Ins1^Cre/+^*; *Dpy30^flox/flox^*; WT: *Ins1^+/+^*; *Dpy30^flox/flox^*; HET: *Ins1^Cre/+^*; *Dpy30^flox/+^*. Mice were maintained on a mixed genetic background. At 8-weeks-old, *Pdx1-CreER* mice were administered 8 mg tamoxifen (Sigma) dissolved in 100 μL corn oil by oral gavage three times, with ∼48-hours between administrations. The first day of injection is considered day 0. BKS *Lepr^db/db^* mice and BKS *Dock7^m/m^* controls (Jax 000642) were used at 12-weeks-old. Procedures were restricted to male mice and approved by the University of British Columbia Animal Care Committee under certificates A17-0045 and A18-0111. Up to five littermates were housed per cage with temperature and humidity control under a 12-hour light/dark cycle and ad libitum access to water and food (Teklad 2918).

### Cell culture

Primary mouse islet cells were maintained in RPMI 1640 medium containing 11 mM glucose (Gibco), 10% (v/v) FBS, 50 U/ml penicillin, 50μg/mL streptomycin (complete RPMI) in 37°C, 5% CO2 atmosphere. *Drosophila* S2 cells were maintained at ambient temperature and air in Schneider’s Drosophila Medium (Gibco) containing 10% (v/v) heat-inactivated FBS, 50 U/ml penicillin, 50μg/mL streptomycin.

### Islet and β-cell isolation

Unless otherwise indicated, mouse islets were isolated 45-days after tamoxifen administration (i.e. at 14- weeks old) and processed for analysis immediately after isolation. Pancreata were perfused with 700 U/mL collagenase type XI (Sigma-Aldrich) in Hank’s Buffered Saline Solution (138 mM NaCl, 5.3 mM KCl, 0.44 mM KH2PO4, 0.34 mM Na2HPO4, 5.5 mM D-Glucose, pH 7.3) through the bile duct, excised, and held for 15-minutes at 40°C before manual disruption and hand-picking of islets. To isolate β-cells, islets from *Pdx1-CreER^Tg/^*^0^ *Rosa26^mTmG/-^* transgenic mice were washed 3× in room temperature PBS + 2 mM EDTA and then dispersed by constant gentle pipetting in 1 mL of 0.025% Trypsin-EDTA (Thermo Fisher Scientific) in PBS at room temperature for 8-10 minutes. Trypsin was quenched by addition of 9 mL room temperature complete RPMI, cells were centrifuged for 3-minutes at 600 × *g* at 4°C, and resuspended in 0.3-0.5 mL ice-cold sorting buffer (PBS, 2 mM EDTA, 1% (w/v) BSA). After passing through a 70 μm cell strainer, singlet eGFP+ tdTomato- cells were purified by fluorescence-activated cell sorting (FACS) using a FACS-Aria II with a 100 μm nozzle.

### Chromatin immunoprecipitation sequencing (ChIP-seq)

A native ChIP protocol ^69^ was used to generate ChIP-seq libraries in biological duplicate, with modifications. For *Pdx1-CreER^Tg/^*^0^; *Dpy30^+/+^*; *Rosa26^mTmG/+^* and *Pdx1-CreER^Tg/^*^0^; *Dpy30^flox/flox^*; *Rosa26^mTmG/+^* mice, 100,000 eGFP+ tdTomato- islet cells were pooled with 50,000 *Drosophila* S2 cells by FACS. For *Lepr^db/db^* and *Lepr^+/+^* mice, 100,000 dispersed islet cells were counted by hemacytometer and pooled with 50,000 *Drosophila* S2 cells without sorting. Pooled cells were centrifuged at 600 × *g* for 5 minutes at 4°C, supernatant removed, and then cell pellets were flash-frozen and stored at -80°C for up to one month. Upon thawing, cells were permeabilized in nuclear isolation buffer (0.1% Triton X-100, 0.1% sodium deoxycholate) containing 1 mM phenylmethylsulfonyl fluoride (PMSF) and EDTA-free protease inhibitor cocktail (Roche)) on ice. Chromatin was fragmented to mononucleosomes by micrococcal nuclease (MNase) (NEB) in digestion buffer (50 mM Tris pH 7.9, 5 mM CaCl2, 1.5 mM DTT) for 7.5-minutes at 37°C, then quenched by addition of 0.1 volume of a solution containing 2% Triton X-100, 2% sodium deoxycholate, and 100 mM EDTA. Soluble chromatin was pre-cleared with Protein A Dynabeads (Thermo Fisher Scientific) pre-adsorbed with normal rabbit IgG (Millipore, 12-370) for 2-hours at 4°C rotating 9 rpm, and then rotated overnight at 4°C with 1 μg of H3K4me3 (Abcam, ab1012), 2 μg H3K4me1 (Abcam, ab8895), 1 μg H3K27ac (Active Motif, 39034), or 2 μg H3K27me3 (Millipore, 07-449) antibodies pre-adsorbed to 10 μL of Protein A Dynabeads (Thermo Fisher Scientific) in 150 μL of ChIP buffer (20 mM Tris pH 8.0, 2 mM EDTA, 150 mM NaCl, 0.2% Triton X-100, 1 mM PMSF, and EDTA-free protease inhibitor cocktail (Roche)). After washing the beads twice with ice-cold wash buffer (20 mM Tris pH 8, 150 mM NaCl, 2 mM EDTA, 1% (v/v) Triton X-100, 0.1% (w/v) SDS) and twice with ice-cold high-salt wash buffer (20 mM Tris pH 8, 500 mM NaCl, 2 mM EDTA, 1% (v/v) Triton X-100, 0.1% (w/v) SDS), DNA was eluted in elution buffer (100 mM NaHCO3, 1% (w/v) SDS, 50 μg/mL RNase A) for 1-hour at 55°C, with addition of 1 μL 20 mg/mL proteinase K after the first 30-minutes and occasional manual inversion. DNA was purified using standard Phenol:chloroform:isoamyl alcohol extraction and ethanol precipitation and used for library preparation with the NEBNext Ultra II DNA Library Prep Kit for Illumina (NEB) with 9 (H3K4me1, H3K27me3) or 13 (H3K4me3, H3K27ac) amplification cycles. Indexed libraries were analyzed for size distribution using the Bioanalyzer High Sensitivity DNA chip (Agilent) and for concentration using the Qubit dsDNA HS Assay (Thermo Fisher Scientific). *Dpy30*-KO samples were sequenced on the NextSeq 500 platform (Illumina) for 2 × 78 nucleotide paired end reads and *Lepr^db/db^* samples on the NextSeq 2000 platform (illumina) for 2 × 61 nucleotide paired end reads. Sequence data is available in the Gene Expression Omnibus (GEO) under accession number GSE181951.

### ChIP-seq data analysis

ChIP-seq reads were aligned to concatenated mouse (GRCm38/mm10) and *Drosophila* (BDGP6.28) genome assemblies using Bowtie2 v2.3.4.1 with options ‘--very-sensitive --no-unal --no-discordant’ ^70^. Multi-mapped reads, reads with mapping quality MAPQ < 20, and suspected PCR-duplicates were discarded using Samtools ^71^. To scale enrichment according to the *Drosophila* cell spike-ins, the fraction of remaining reads that mapped to the *Drosophila* genome was determined, and a scaling factor was calculated for each sample as *(minimum* Drosophila *read fraction per histone mark)/(sample* Drosophila *read fraction)*. After determining a scaling factor, each sample was downsampled to the lowest mapped read count per each histone mark using Samtools, and then reads mapped to the *Drosophila* genome were removed. For visualization, libraries were converted to bedgraph format using the calculated scaling factor in the ‘-scale’ argument of BEDtools v2.26.0 genomecov ^72^. Genome browser views were generated for merged replicates using Spark ^73^ and TSS profiles using Deeptools v3.4.2 ^74^. Peak locations and breadth were determined using MACS2 v2.2.6 with parameters ‘--broad --nolambda’ ^41^. Peaks detected in only one biological replicate or that overlapped with high-background blacklisted regions defined by the ENCODE project ^75^ were excluded. Chromosomes X, Y, and M were also excluded from the analyses. DESeq2 ^76^ was used for differential peak width analysis using a significance cutoff of *P* ≤ 0.05 (Wald test with Benjamini and Hochberg adjustment), and enrichment for differential breadth in peak quantiles was determined by one-sided Fisher’s exact test. We defined TSS locations as the 5’ end of the most abundant annotated transcript of each gene detected in our RNA-seq data. H3K4me3 peaks were assigned to genes for which they overlap the region ±1 kb of the TSS.

### RNA-seq

eGFP+ tdTomato- β-cells (70,000-226,000 per mouse, biological triplicate) were purified and counted by FACS and supplemented with a 10% spike-in of *Drosophila* S2 cells. *Drosophila* cells were included to detect global changes in cell RNA content ^77^. The cells were sorted directly into Trizol LS Reagent (Thermo Fisher Scientific) and total RNA was extracted according to manufacturer’s instructions. Residual DNA was digested by Turbo DNase (Thermo Fisher Scientific) for 30-minutes at 37°C. Purified total RNA content was measured using the Qubit RNA HS assay (Thermo Fisher Scientific). Mature mRNA was enriched from 400 ng of total RNA per sample using the NEBNext Poly(A) mRNA Magnetic Isolation Module (NEB) and used to prepare libraries with the NEBNext Ultra II Directional RNA Library Prep Kit for Illumina (NEB) with 11 amplification cycles. Uniquely indexed libraries were quantified, pooled, and sequenced for 2 × 38 nucleotide reads as described above.

### RNA-seq data analysis

Sequenced reads were aligned to the concatenated mouse (GRCm38/mm10; Gencode vM24) and *Drosophila* (BDGP6.28; Ensembl release 99) genomes and transcriptomes using STAR v2.7.3a ^78^ with default settings. Transcript abundance was calculated using Salmon v1.4.0 ^79^ in alignment mode with seqBias and gcBias options. Gene counts and differential expression were calculated using DESeq2 ^76^. Genes with fewer than two counts in any sample were filtered out and genes showing ≥ 2-fold difference in expression with *P* ≤ 0.01 (Wald test with Benjamini and Hochberg adjustment for multiple comparisons) are considered differentially expressed. Genes showing <1.1 fold difference in expression are considered stable. Except in Fig. 6F, Drosophila transcripts were removed after abundance calculation; for Fig. 6F, *Drosophila* transcripts were used as control genes in the estimateSizeFactors function of DESeq2, and then removed.

### Single-cell RNA-seq

Islets from one WT and one KO mouse were dissociated to single-cell suspensions as described above without FACS-enrichment. The cell suspensions were processed through the Chromium Single Cell 3’ protocol using the Chromium Controller (firmware v4.0) with Reagent Kit v3.1 (10X Genomics) and Dual Index Kit TT Set A (10X Genomics) according to the manufacturer’s instructions for a targeted 5,000-cell recovery. Total cDNA was amplified for 11-cycles, and then 13-cycles during index-ligation. cDNA concentrations were quantified by qPCR with the NEBNext Library Quant Kit for Illumina (NEB). Libraries were pooled and sequenced for 28-90 paired-end nucleotide reads to an average depth of ∼70,000 mapped fragments per cell.

### Single cell RNA-seq data analysis

Cellranger (v5.0.0, 10X Genomics) was used to generate FASTQ files and demultiplex reads. Cell barcode detection, read mapping, quality filtering, and transcript counting were performed using Alevin (Salmon v1.4.0) ^80^ against the mouse protein-coding transcriptome (Gencode vM24), which we appended with the *Tdtomato* and *Egfp* sequences, as well as the mouse genome (GRCm38/mm10) as mapping decoys for selective alignment ^81^. Additional cell quality filtering was performed using Seurat v4.0 ^82^ based on unique features (> 1,000 genes) and mitochondrial RNA content (< 25%). Putative doublets were excluded using DoubletFilter ^83^. Genes detected in < 3 cells were excluded from analysis. After quality filtering, the dataset comprised 3,703 WT and 3,641 KO cells. Filtered data were scaled and normalized using SCTransform ^84^ with default parameters. PCA and UMAP dimensionality reduction using the first 60 principal components was used to cluster and visualize cell populations. We then used Slingshot ^85^ to perform pseudotime analysis on β-cell clusters. Entropy scores are a modified Shannon entropy calculated using a published R function (ref ^86^).

### Immunohistochemistry

Mouse pancreata were isolated and fixed in 4% (w/v) paraformaldehyde (PFA) in PBS at 4°C overnight. Following washing with PBS and dehydration in ethanol and xylenes, mouse tissues were embedded in paraffin and sectioned with a thickness of 5 μm. Immunofluorescent analyses of tissue sections was performed as previously described^23^ in a blinded fashion from 4-6 sections per biological sample. Primary antibodies used were: H3K4me3 (Cell Signaling Technologies, C42D8, 1:500), H3K4me1 (Abcam, ab8895, 1:1,000), H3 (Abcam, ab1791, 1:1,000), DPY30 (Atlas Antibodies, HPA043761, 1:500), Insulin (Agilent, IR-002, 1:4), and Glucagon (Sigma Aldrich, G2654, 1:1,000).

### Transmission electron microscopy

Islets were fixed in 2% glutaraldehyde (Sigma Aldrich) in PBS and submitted to the Facility for Electron Microscopy Research of McGill University (Hamilton, ON, Canada). Islets were post-fixed with 1% osmium tetroxide, dehydrated in ethanol, and embedded in Spurr’s resin before sectioning with a Leica UCT ultramicrotome. Sections were stained with uranyl acetate and lead citrate and imaged with a JEOL JEM 1200 EX TEMSCAN transmission electron microscope (JEOL, Peabody, MA, USA).

### Immunoblotting

Islets were lysed in Laemmli sample buffer (2% (w/v) SDS, 10% (v/v) glycerol, 60 mM Tris pH 6.8, 1mM NaF, 1 mM PMSF, protease inhibitor cocktail (Roche)) for 5-minutes at 95°C. Protein concentration was measured by BCA assay (Thermo Fisher Scientific), then β-mercaptoethanol was added to 5% (v/v) and lysates were reboiled for 5-minutes and stored at -80°C. 10 μg lysates were resolved in polyacrylamide gels, transferred to PVDF membranes, blocked for 60-minutes in 5% (w/v) skim milk powder in TBST (20 mM Tris pH 7.6, 150 mM NaCl, 0.1% (v/v) Tween-20) at room temperature and probed with primary antibodies overnight in 5% (w/v) BSA in TBST at 4°C. The next day, after washing membranes three times with TBST, HRP-linked secondary antibodies were added for 1-hour at room temperature, membranes were washed three times with TBST, and detected using ECL system with film. When normalizing to total H3, membranes were stripped for 20-minutes in mild stripping buffer (200 mM glycine, 0.1% (w/v) SDS, 1% (v/v) Tween-20, pH 2.2) at room temperature, rinsed twice in PBS and twice in TBST for 10 minutes each at room temperature, then re-blocked and probed for histone H3. Primary antibodies used for immunoblots were: H3K4me3 (Cell Signaling Technologies, C42D8, 1:1,000), H3K4me1 (Abcam, ab8895, 1:100,000), H3K27ac (Cell Signaling Technologies, D5E4, 1:1,000), H3K27me3 (Millipore, 07-449, 1:100,000), H3 (Abcam, ab1791, 1:1,000,000), WDR5 (Bethyl, A302-430A, 1:2,000), RBBP5 (Bethyl, A300-109A, 1:10,000), and ASH2L (Bethyl, A300-489A, 1:10,000).

### Protein co-immunoprecipitation

Immunoprecipitation of WDR5 was performed in triplicate in 750-900 dispersed islets pooled from 3 mice. First, nuclei were enriched from dispersed islet cells by incubation in nuclear isolation buffer for 10- minutes on ice and pelleting at 600 × *g* for 5-minutes at 4°C. Chromatin was solubilized in IP buffer (20 mM Tris pH 7.4, 150 mM NaCl, 1 mM EDTA, 5% (v/v) glycerol, 0.5% (v/v) Igepal ca-630, 5 mM CaCl2, 1 mM PMSF, protease inhibitor cocktail (Roche)) with MNase (NEB) for 10-minutes at room temperature. EDTA was added to 5 mM to stop MNase and samples were centrifuged at 14,000 × *g* for 5-minutes at 4°C. The protein concentration in the supernatant was measured by BCA assay (Thermo Fisher Scientific), and an equal mass of WT and KO nuclear lysate (∼30 μg protein) was immunoprecipitated by 1 μg WDR5 (Bethyl, A302-429A) or normal rabbit IgG (Millipore, 12-370) antibodies rotating overnight at 4°C. The next day, 10 μL protein A Dynabeads (Thermo Fisher Scientific) was added to the antibody:lysate mixture and rotated for 2-hours at 4°C. Finally, bead:antibody:antigen complexes were washed six times in ice-cold IP buffer and resuspended in Laemmli sample buffer with 5% (v/v) β-mercaptoethanol for immunoblotting.

### Insulin secretion assays

For in vivo glucose tolerance tests, 2 g/kg glucose in a 20% (w/v) water solution was injected intraperitoneally following a 6-hour fast. Blood glucose was sampled immediately prior to and 15, 30, 60, and 120-minutes after glucose injection from a lateral saphenous vein. Serum was collected at the 0, 15, and 60-minute time points by centrifuging blood samples at 9,000 × g for 9 minutes at 4°C and stored at -80°C. Serum insulin was measured by ELISA (Alpco) according to the manufacturer’s instructions. Ex vivo static insulin secretion assays were performed in technical duplicate in groups of 50 islets per mouse one day after isolation. Islets were preincubated in KRBH buffer (2.8 mM glucose, 20 mM HEPES, 114 mM NaCl, 4.7 mM KCl, 1.2 mM MgSO4, 1.2 mM KH2PO4, 2.5 mM CaCl2, 24 mM NaHCO3, 0.1% (w/v) BSA, pH 7.3) for one hour at 37°C and then sequentially for 45-minutes in KRBH, either 16.7 mM glucose or 15 mM dimethyl α-ketoglutarate (Sigma Aldrich), and finally 30 mM KCl. Conditioned medium was collected after each incubation. At the completion of the assay islets were lysed in RIPA buffer (Pierce) containing 1 mM PMSF and EDTA-free protease inhibitor cocktail (Roche)) at 95°C for 5 minutes. Insulin concentration was measured in conditioned medium and islet lysate by ELISA (Alpco). DNA concentration of islet lysate was determined by Qubit dsDNA HS Assay (Thermo Fisher Scientific) to normalize islet insulin content.

### Calcium imaging

Intact islets were cultured for two days to allow for attachment onto glass coverslips. Prior to imaging, coverslips were transferred to a 2 mL imaging chamber and stained with 5 μM Fura-2AM (Molecular Probes, F1221) for 30-minutes at 37°C in complete RPMI. The imaging chamber loaded with a coverslip was then mounted on a Leica SP8 Laser Scanning Confocal Microscope with a 10× objective lens and continuously washed with Ringer’s solution (5.5 mM KCl, 2 mM CaCl2, 1 mM MgCl2, 20 mM HEPES, 144 mM NaCl, 2.8 mM glucose, pH 7.4) for 30-minutes. Throughout the experiment, islets were continuously perifused at a flow rate of 2.5 mL/min with Ringer’s solution. Glucose and KCl concentrations were adjusted by iso-osmotic substitution of NaCl where indicated. Fura-2 was excited at 340 nm and 380 nm and emitted fluorescence was detected at approximately 510 nm. Cytosolic Ca^2+^ levels are expressed as the mean ratio of Fura-2 fluorescence emission intensity (F340/F380) of 3-9 islets per biological replicate.

### Respirometry

Islets were dissociated to single cells 3-4 hours after isolation and plated at 100 islets/well in a Seahorse 24-well cell culture dish in technical duplicate. Cells were cultured for 2-days in complete RPMI to allow for attachment to the culture dish and then for 1-hour in XF RPMI medium (Agilent) with 2.8 mM glucose, 1% FBS, 1 mM pyruvate, and 2 mM glutamine to reach a metabolic baseline. Oxygen consumption was then measured using a Seahorse XF24 Extracellular Flux Analyser (Agilent). The injection ports were loaded with the following compounds in XF Assay Media (Agilent): A, 167 mM glucose; B, 10 μM oligomycin; C, 15 μM FCCP; D, 10 μM antimycin A and 1 μM rotenone. Measurements were normalized to the DNA content of assayed cells, which were determined using Qubit HS DNA Assay (Thermo Fisher Scientific) after lysing in RIPA buffer upon completion of the assay.

### Analysis of publicly available data

Publicly available data were reprocessed for consistency. GEO: GSE50244, GSE50386, GSE107489, GSE124742 were downloaded from the Sequence Read Archive (SRA) using the sratoolkit (https://github.com/ncbi/sra-tools) and analyzed according to the experimental application, as described above.

### Statistics

Unless otherwise stated, plots show mean ± standard deviation (SD) with points representing each biological replicate. In boxplots, the central horizontal line marks median, the upper and lower limits of the box the first and third quartiles, and whiskers span 1.5× the interquartile range, in the style of Tukey. Spearman’s rank coefficient was used to estimate correlation. Regression lines were generated using a generalized additive model. *P*-values were calculated using Student’s t-tests with Welch’s correction, Wilcoxon rank sum test, Wald test, mixed-effect model, one- or two-way ANOVA, Fisher’s exact test, or permutation test, as indicated. Correction for multiple comparison using the Benjamini-Hochberg method was applied where indicated. Statistical calculations were performed in R or Prism 8 or 9 (GraphPad Software, La Jolla, CA, USA). *P* < 0.05 was deemed significant.

## Author contributions

Experiments were conducted by B.V., N.M., D.J.P., M.A., and C.L.M, and conceptualized by B.V. and B.G.H. D.J.P. and D.S.L. performed calcium imaging experiments and designed respirometry experiments. Bioinformatics were performed by B.V. and B.G.H. Funding and supervision was provided by B.G.H. and F.C.L; the manuscript was written by B.V. B.G.H. is the guarantor of this work and, as such, had full access to all the data in the study and takes responsibility for the integrity of the data and the accuracy of the data analysis.

## Acknowledgements

The authors thank the staff of the Animal Care Facility at British Columbia Children’s Hospital Research Institute for daily maintenance of the mouse colonies, and W. Gibson (UBC) for providing Ins1^Cre^ animals. We thank M. Reid at the Facility for Electron Microscopy Research of McGill University for help with transmission electron microscopy, L. Xu at the Flow Core of British Columbia Children’s Hospital Research Institute for help with FACS. This work was supported by operating grants from the British Columbia Children’s Hospital Research Institute, the Natural Sciences and Engineering Research Council of Canada (RGPIN-2016-04292) and the Canadian Institute for Health Research (RN310864 – 375894). Fellowship support for B.V. was provided by the British Columbia Children’s Hospital Research Institute, University of British Columbia, and Natural Sciences and Engineering Research Council of Canada.

---

**Figure S1.**
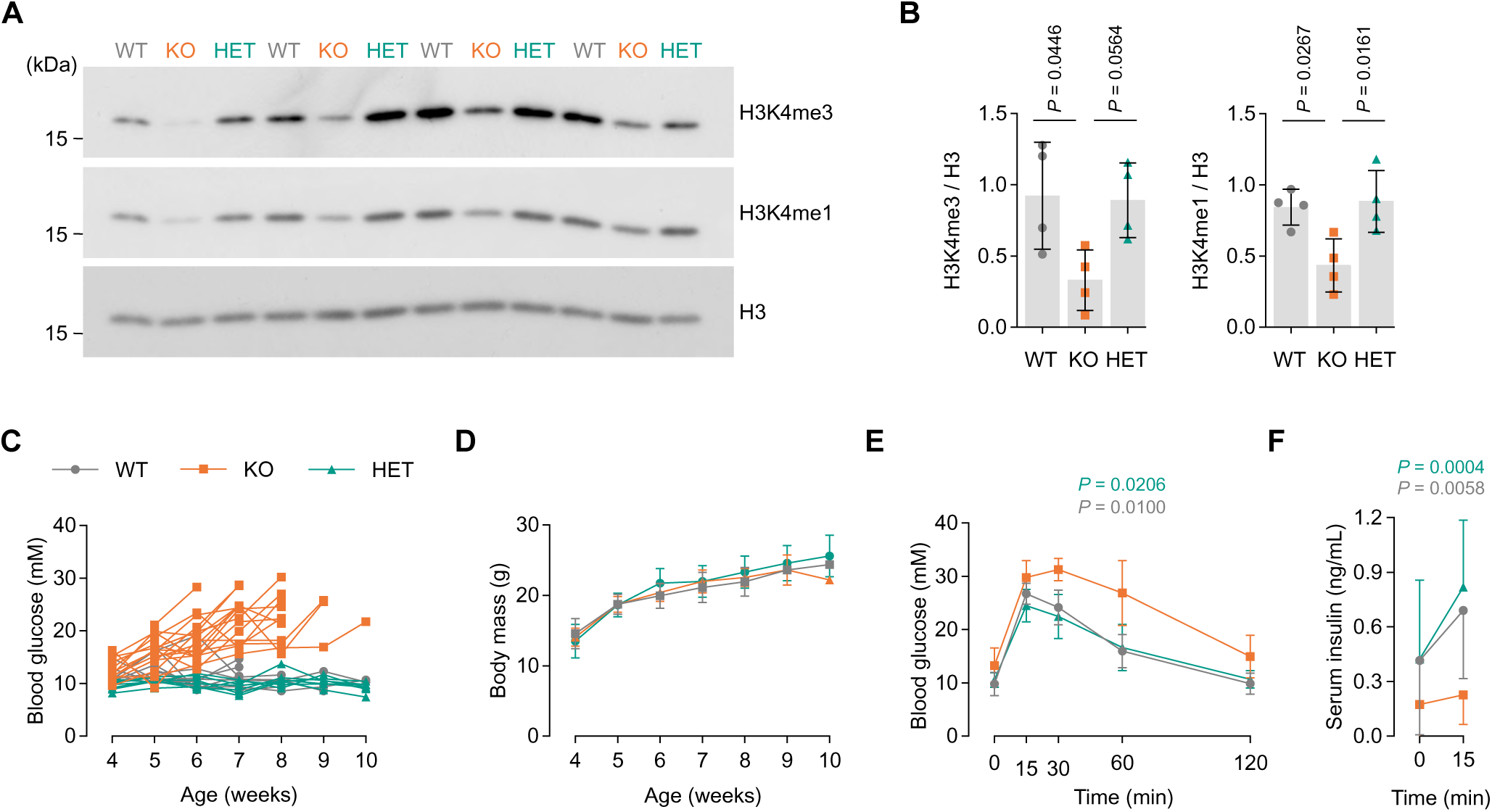
Deletion of *Dpy30* in mouse β-cells using *Ins1^Cre^* results in reduction of H3K4 methylation, hyperglycemia, hypoinsulinemia, and impaired glucose tolerance. A, Immunoblots of H3K4me3, H3K4me1, and total histone H3 in islet lysate from 5-week-old *Dpy30*-WT, -KO, and -HET mice. **B,** Bar graphs showing band intensities of data in panel A. *P*-values were calculated using one-way ANOVA with Tukey’s multiple comparisons correction; n = 4. **C-D,** Unfasted blood glucose concentration (C) and body mass (D) of *Dpy30*-WT, -KO, and -HET mice during 4-10 weeks of age. Values from each mouse and time point are shown (n = 10 WT, 15 KO, 7 HET, however tracking was stopped after 1-2 blood glucose reading ≥ 20 mM). **E-F,** Blood glucose (E) and serum insulin (F) concentration during intraperitoneal glucose tolerance tests in 5-week-old *Dpy30*-WT, -KO, and -HET mice. *P*-values were calculated by comparison of *Dpy30*-KO versus -HET (top, teal) or -KO versus -WT (bottom, grey) AUC’s using one-way ANOVA with Tukey’s multiple comparisons correction; only values ≤ 0.05 are shown. WT: *Ins1^+/+^*; *Dpy30^flox/flox^*. KO: *Ins1^Cre/+^*; *Dpy30^flox/flox^*. HET: *Ins1^Cre/+^*; *Dpy30^flox/+^*.

**Figure S2.**
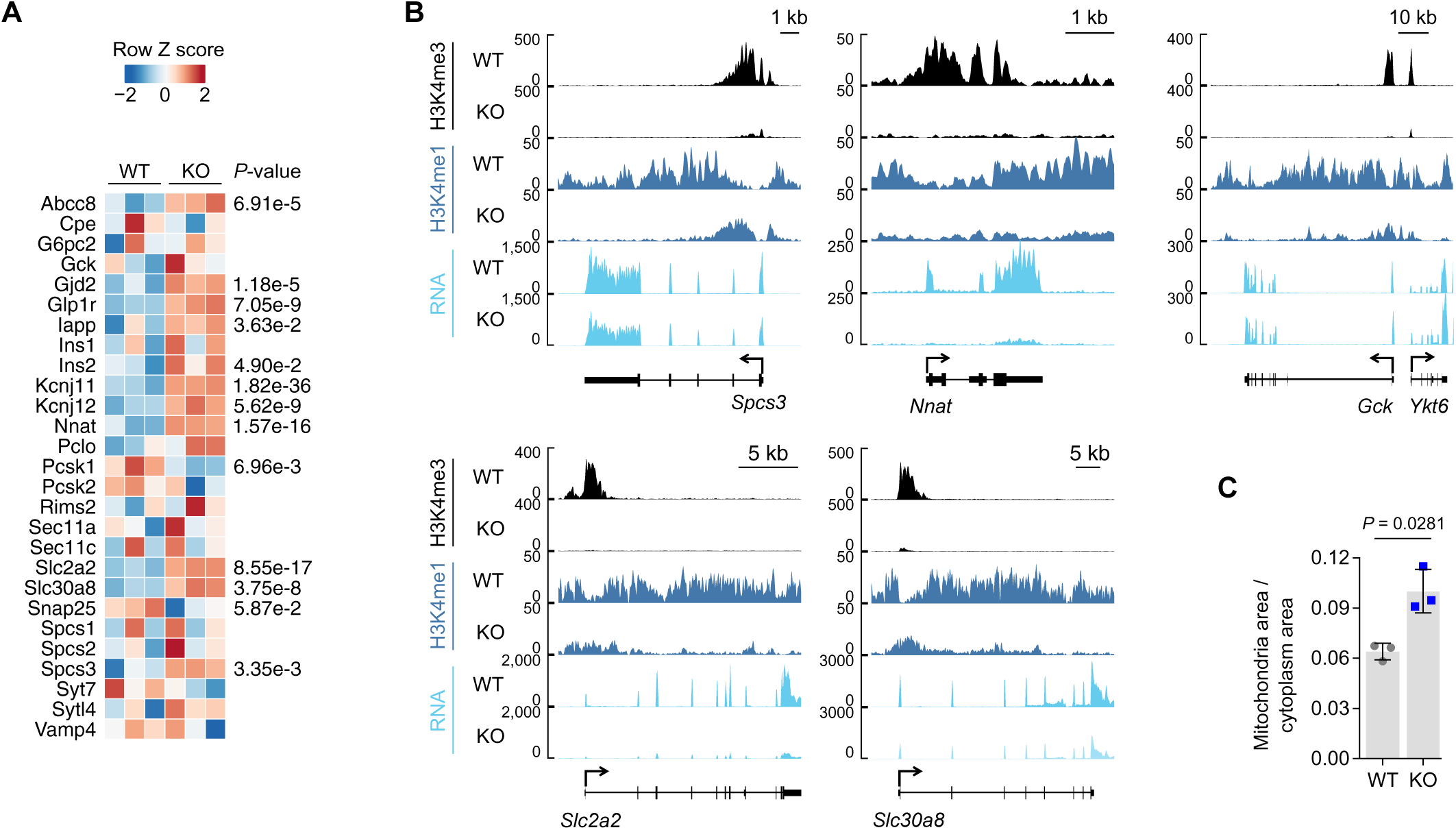
Genes involved in insulin production and glucose-induced activity are regulated by H3K4me3. **A,** Heatmap showing expression Z-scores in RNA-seq data from *Dpy30*-WT and -KO 45-days post-tamoxifen for selected genes important for glucose metabolism, insulin production or secretion. *P*-values calculated by Wald test with Benjamini-Hochberg correction; n = 3. **B,** Genome browser views of H3K4me3, H3K4me1, and RNA in *Dpy30*-WT and -KO chromatin at selected genes. **C,** Bar graph showing the fraction of cytoplasm area occupied by mitochondria in transmission electron micrographs of *Dpy30*- WT and -KO β-cells. *P*-value calculated using two-tailed t-test with Welch’s correction. **D,** Representative immunofluorescence images showing EdU incorporation in *Dpy30*-WT and -KO mouse islets that were dispersed and cultured in 5.5 mM or 16.7 mM glucose for 48-hours, as indicated, showing EdU (white) and DAPI (white) staining and eGFP (green) and tdTomato (red) endogenous fluorescence. Scale bars: 100 μm. **E,** Bar graph showing quantification of EdU+ eGFP+ double positive cells as a fraction of total eGFP+ cells (n = 3 per treatment and genotype). *P*-values calculated using two-way ANOVA with Tukey’s correction. ns: not significant (*P* > 0.05).

**Figure S3.**
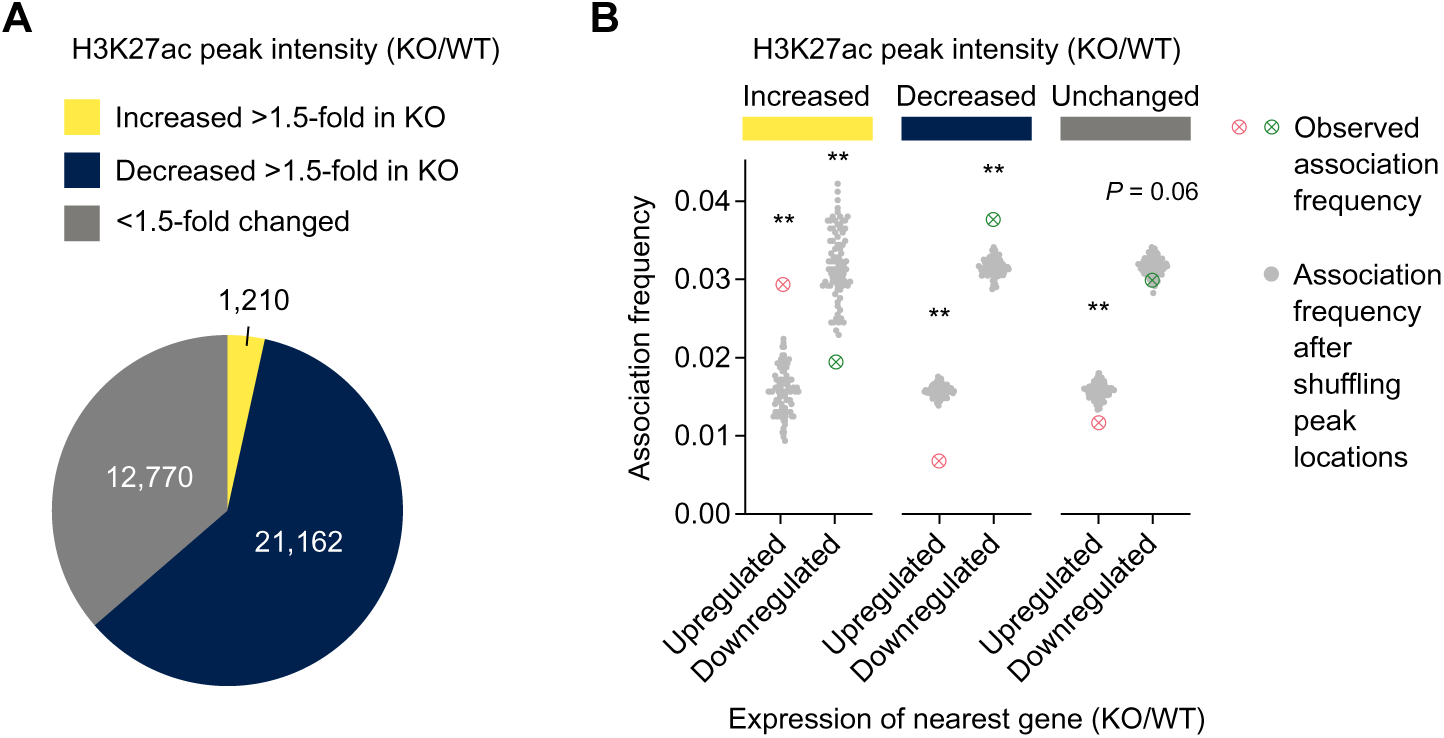
H3K27ac peak dynamics are linked to gene expression dynamics in *Dpy30*-KO cells. **A,** Pie chart showing the number of H3K27ac peaks that are increased >1.5-fold, decreased >1.5-fold, or <1.5-fold changed for enrichment of H3K27ac in *Dpy30*-KO versus -WT chromatin. **B,** Association frequency of H3K27ac peaks classified in panel A with genes that are upregulated or downregulated in *Dpy30*-KO cells. Association was tested between H3K27ac peaks with the nearest active TSS and compared with association frequencies after randomizing peak locations 100 times. ** *P* < 0.01 by permutation test.

**Figure S4.**
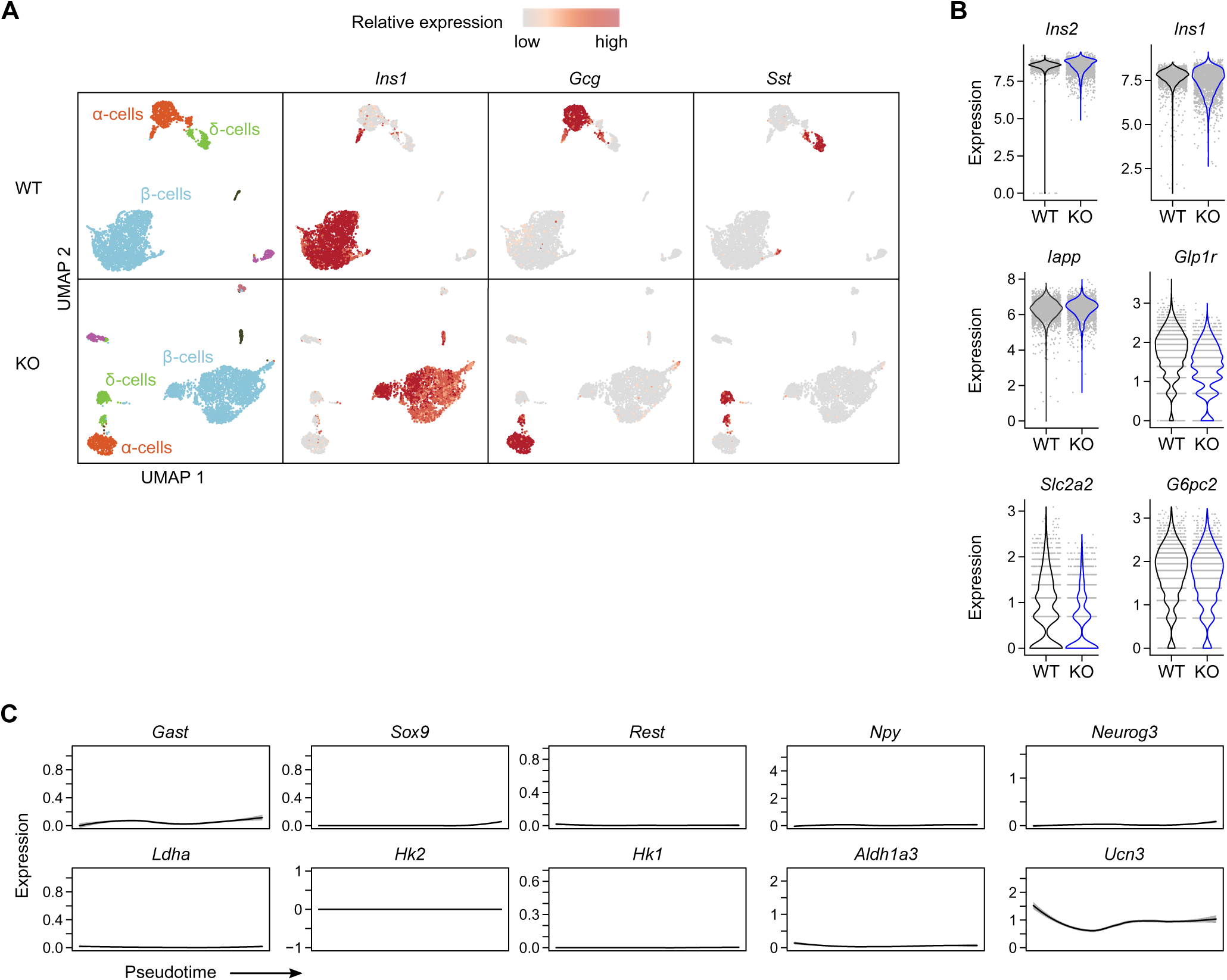
*Dpy30*-KO β-cells do not activate genes associated with dedifferentiated β-cells. **A,** UMAP visualization of islet cell transcriptomes from a *Dpy30*-WT and -KO mouse. Markers of β-cells (*Ins1*), α-cells (*Gcg*), and δ-cells (*Sst*) are shown as a colour gradient (n = 3,703 WT cells, 3,641 KO cells). **B,** Violin plots showing expression *Ins1*, *Ins2*, *Iapp*, *Glp1r*, *Slc2a2*, and *G6pc2* in *Dpy30*-WT and -KO β-cells (n = 2,510 WT, 2,597 KO β-cells). **C,** Lowess curves of gene expression of the indicated maturity and immaturity genes in *Dpy30*-KO β-cells along pseudotime defined in Fig 6B.

**Figure S5.**
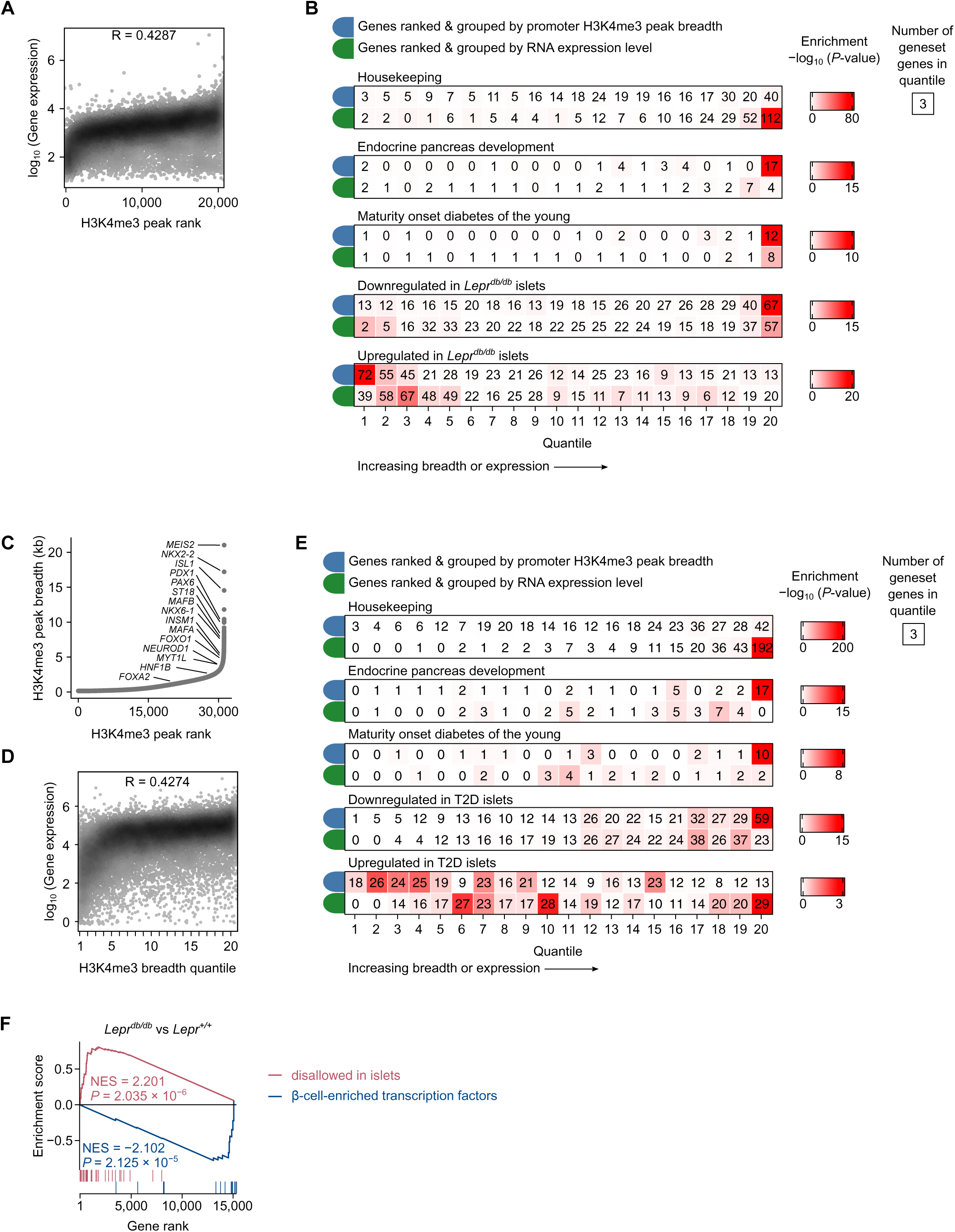
H3K4me3 peak breadth stratifies β-cell gene sets more effectively than gene expression level. **A,** H3K4me3 peaks ranked by increasing breadth (x-axis) plotted against RNA expression of the associated gene (y-axis). R indicates Spearman’s rank correlation coefficient. **B,** Overrepresentation analysis of expressed genes that are ranked and grouped by H3K4me3 peak breadth from narrow to broad (blue) or ranked and grouped by gene expression level from low to high expression (green). The number in each square indicates the number of genes of each geneset matching to each quantile. Enrichment *P*-values calculated using two-sided Fisher’s exact tests. **C,** H3K4me3 peaks in human β-cells ^51^ ranked by peak breadth. Peaks associated with β-cell transcription factor genes are labeled. **D,** H3K4me3 peaks ranked by breadth (x-axis) plotted against RNA expression ^58^ of the associated gene (y-axis) in non-diabetic human islets. R indicates Spearman’s rank correlation coefficient. **E,** Overrepresentation analysis of human islet genes that are ranked and grouped by H3K4me3 peak breadth from narrow to broad (blue) or ranked and grouped by gene expression level from low to high expression (green). The number in each square indicates the number of genes of each gene set matching to each quantile. Each quantile includes 783 genes. Enrichment *P*-values calculated using one-sided Fisher’s exact test. **F,** Running enrichment plot of genes disallowed in islets ^54^ and β-cell-enriched transcription factor genes in *Lepr^db/db^* RNA-seq data ^56^ . NES: normalized enrichment score. *P*-values calculated by permutation test.

